# Triosephosphate export from chloroplasts regulates flavonoid biosynthesis and permits high light acclimation through the inactivation of SnRK1

**DOI:** 10.1101/2022.03.09.483619

**Authors:** Max-Emanuel Zirngibl, Galileo Estopare Araguirang, Anastasia Kitashova, Kathrin Jahnke, Tobias Rolka, Christine Kühn, Thomas Nägele, Andreas S. Richter

## Abstract

Plants evolved multiple strategies to cope with rapid changes in the environment. During high light acclimation, biosynthesis of photoprotective flavonoids, such as anthocyanins, is induced. However, the exact nature of the signal and downstream factors for high light induction of flavonoid biosynthesis (FB) are still under debate. Here we show that carbon-fixation in chloroplasts, subsequent export of photosynthates by TRIOSEPHOSPHATE/PHOSPHATE TRANSLOCATOR (TPT), and the rapid increase in cellular sugar contents permit the transcriptional activation of FB during high light acclimation. In combination with genetic and physiological analysis, targeted and whole transcriptome gene expression studies showed that reactive oxygen species and phytohormones play only a minor role for rapid HL-induction of the anthocyanin branch of FB. In addition to FB, sugar-responsive genes were late-repressed or induced in *tpt-2* in the course of the high light treatment and a significant overlap with transcripts regulated by SNF1-RELATED PROTEIN KINASE 1 (SnRK1) was found. Analysis of mutants with increased and repressed SnRK1 activity revealed that inactivation of SnRK1 is required for the rapid induction of FB during high light acclimation. Our study underlines the central role of chloroplasts as sensors for environmental changes and emphasizes the vital function of sugar-signalling in plant acclimation, even beyond the regulation of FB.

## Introduction

Due to their sessile lifestyle, plants evolved mechanisms to adjust cellular processes and metabolisms to cope with sudden environmental changes. Among them, light intensity or temperature variations can occur within seconds to minutes and can persist for hours or several days during the different seasons of a year. Besides rapid adjustments on the metabolic and post-translational level, tuning the gene expression activity is vital for the acclimation response, which depends on substantial transcriptome reprogramming (for example, Alsharafa et al., 2014, Glasser et al., 2014, Calixto et al., 2018, Huang et al., 2019, Garcia-Molina et al., 2020). During the last decades, it became apparent that chloroplasts function as sensors of changes in the environment and emitters of signals necessary for establishing a new cellular homeostasis when plants face adverse conditions (Nott et al., 2006, Pogson et al., 2008, Hausler et al., 2014, Chan et al., 2016, Dietz et al., 2016, Kleine and Leister, 2016, de Souza et al., 2017, Kleine et al., 2021, Schwenkert et al., 2022). Reactive oxygen species (ROS) (Foyer and Noctor, 2015, Mullineaux et al., 2018, Leister, 2019), oxidative cleavage products of carotenoids (Ramel et al., 2012), plastid-derived phytohormones, or changes in the redox status of the plastoquinone pool (Bräutigam et al., 2009, Pesaresi et al., 2009, Petrillo et al., 2014) serve as signals to adjust nuclear gene expression. In addition, metabolic signals, like an intermediate of the plastid-localized methylerythritol phosphate pathway (Xiao et al., 2012), triosephosphate (Vogel et al., 2014, Weise et al., 2019) and other chloroplast stress signals (e.g.,3’-phosphoadenosine 5’-phosphate; PAP, Estavillo et al., 2011) alter the expression of (stress-specific) nuclear genes.

One common trait of plants acclimating to high light (HL) is the accumulation of (coloured) flavonoids. The precursors of these secondary metabolites (phenylpropanoids) and flavonoids themselves play a protective role by absorbing excessive amounts of (UV-) light and by providing high antioxidant capacity to the cell in adverse environmental conditions (Li et al., 1993, Harvaux and Kloppstech, 2001, Gould, 2004, Huang et al., 2010, Emiliani et al., 2013, Xu et al., 2017, Gould et al., 2018, Agati et al., 2020, Agati et al., 2021). Flavonoids, including anthocyanins, are end products of a combined pathway of cytosolic phenylpropanoid and flavonoid biosynthesis (FB), which produce a great diversity of polyphenolic plant secondary metabolites (Vogt, 2010, Petrussa et al., 2013). Plastid-derived phenylalanine is converted to *p-*coumaroyl CoA through PHENYLALANINE AMMONIA LYASE (PAL), CINNAMIC ACID-4-HYDROXYLASE (C4H) and 4-COUMAROYL COA LIGASE (4CL). Subsequently, CHALCONE SYNTHASE (CHS) catalyzes the initial step of FB providing chalcones. Within the next steps of FB, precursor(s) for various flavonoid derivatives, such as (dihydro)flavonols, are synthesized (Fig. 1A). At a branch point of FB, DIHYDROFLAVONOL 4-REDUCTASE (DFR) and LEUCOANTHOCYANIDIN DIOXYGENASE (LDOX) initiate the route towards anthocyanin biosynthesis (Appelhagen et al., 2014). After the synthesis of anthocyanidins, the aglycones of anthocyanins, glycosyl-, methyl- and acyltransferases contribute to the ‘decoration’ of anthocyanins and are responsible for the diversity of anthocyanins found in nature.

**Figure 1:**
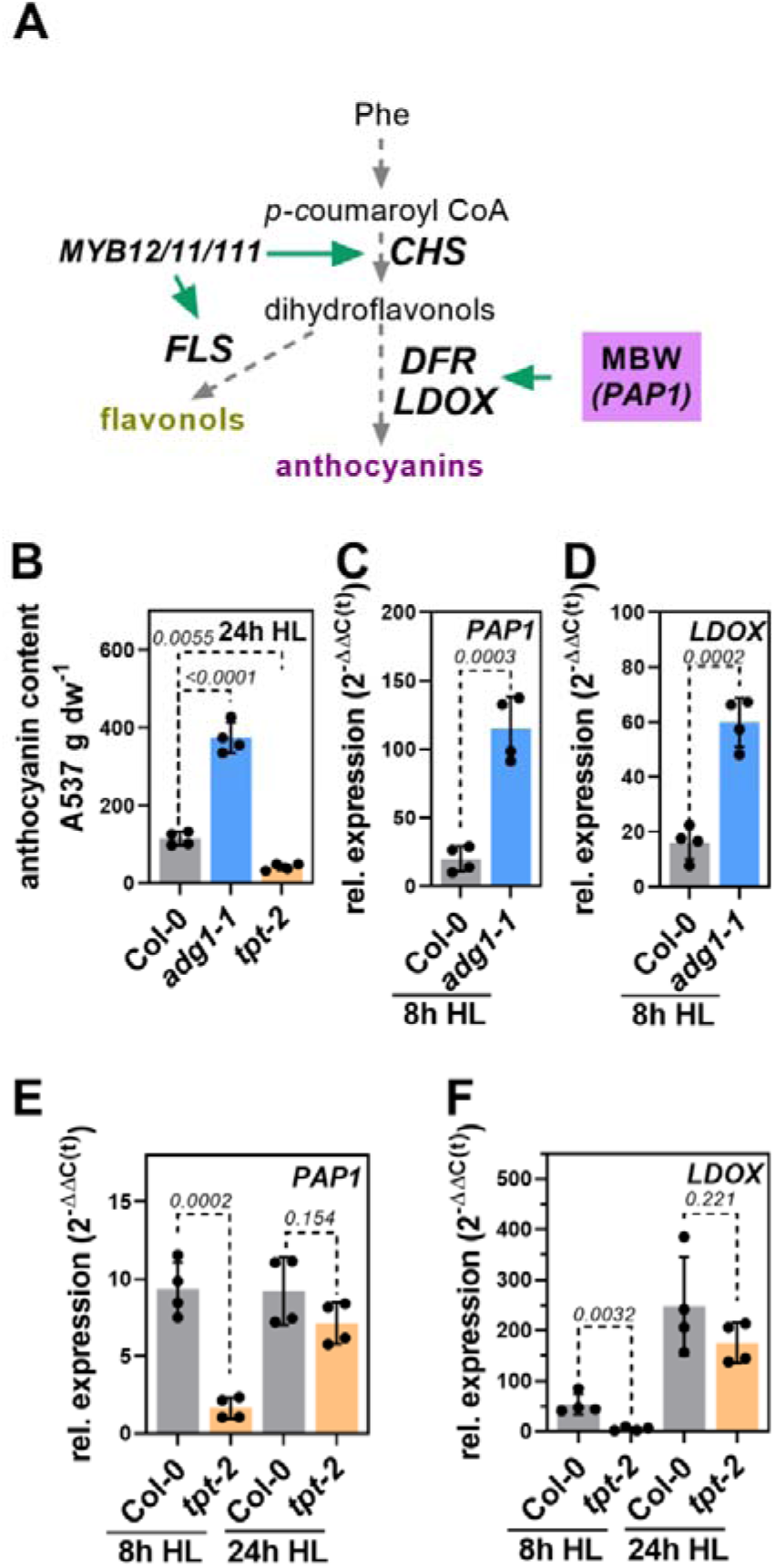
Flavonoid biosynthesis is differentially regulated in starch-deficient *adg1-1* and triose phosphate/phosphate translocator mutant (*tpt-2*). (**A**) Schematic presentation of the biochemical pathway leading to the formation of flavonoids. Further information is provided in the main text. (**B**) Anthocyanin accumulation in wild-type (Col-0), *adg1-1* and *tpt-2* mutant after exposure to continuous high light (HL) for 24h. Plants were shifted to the HL 2h after the onset of light. Statistical significance between wild-type and mutants was calculated using one-way ANOVA (Dunnett’s multiple comparisons test) relative to Col-0. Values are mean ±SD (n=4) and the adjusted *p*-values are shown. (**C-F**) Relative expression of *PAP1* (C, E) and *LDOX* (D, F) in *adg1-1* (C,D) and *tpt-2* (E, F) after 8h HL treatment (*adg1-1* and *tpt-2*) and 24h HL (*tpt-2*). Changes in gene expression were calculated using the 2^−ΔΔ(C(t))^ method and *SAND* as reference gene and are expressed relative to Col-0 before the HL shift. Statistical significance between Col-0 and mutant after 8h and 24h of HL was analyzed using Student’s t-test. Values are mean ±SD (n=4) and the *p*-values are shown.

The FB pathway enzymes/genes are grouped into ‘early biosynthetic genes’ (*EBG*), like *CHS* and *FLAVONOL SYNTHASE1* (*FLS1*), and the ‘late biosynthetic genes’ (*LBG*), such as *DFR* and *LDOX* (Fig. 1). Expression of *EBG* is mostly under the control of the partially redundant transcription factors (TFs) *MYB11, MYB12*, and *MYB111* (Stracke et al., 2007). In contrast, the expression of *LBG* mainly depends on an MBW-complex formed by <u>M</u>YB, <u>b</u>HLH, and <u>W</u>D40 components. Together with one WD40 variant (*TRANSPARENT TESTA GLABRA 1, TTG1*), a group of MYB TFs and the bHLH TFs *TRANSPARENT TESTA 8* (*TT8*), *GLABRA3* (*GL3*) and *ENHANCER OF GL3* (*EGL3*) regulate the composition and activity of the MBW-complex and the overall activity of the FB pathway (Zhang et al., 2003, Lloyd et al., 2017, LaFountain and Yuan, 2021). MYB components acting as positive regulators of FB are encoded by *MYB75* (*PRODUCTION OF ANTHOCYANIN PIGMENT 1, PAP1*), *MYB90* (*PAP2*) and *MYB113* and *114* (Borevitz et al., 2000, Gonzalez et al., 2008). Knock-out of *PAP1* abolishes the expression of FB pathway genes and anthocyanin accumulation (Teng et al., 2005, Li et al., 2016) and is therefore considered to be the major MYB component of the MBW-complex driving FB gene expression. Transcriptional induction of positive regulators precedes the stimulation of pathway genes and results in anthocyanin accumulation under various (stress) conditions. *LBG* are also repressed by the concurred interaction between stimulating and repressive MYBs with the MBW-complex, such as *MYB-LIKE 2* (Dubos et al., 2008, LaFountain and Yuan, 2021).

Enzymes and regulators of FB are targets for post-translational protein modifications, particularly ubiquitination and subsequent proteasomal degradation (Maier et al., 2013, Zhang et al., 2017). In this context, FB genes and TFs are under the control of light-signalling pathways involving UV-, blue (CRYPTOCHROMES, CRY), and red-light (PHYTOCHROMES) photoreceptors and the downstream component ELONGATED HYPOCOTYL 5 (HY5) (Kleine et al., 2007, Stracke et al., 2010, Warnasooriya et al., 2011, Heijde et al., 2013, Shin et al., 2013, Maier and Hoecker, 2015, Job et al., 2018, Ponnu et al., 2019, Bursch et al., 2020). Also, various plastid-derived signals or signals depending on chloroplast function exhibit a function in FB regulation. For example, ascorbate or phytohormones, such as jasmonate, gibberellic acid, auxin, abscisic acid and ethylene, are associated with the regulation of FB (Dombrecht et al., 2007, Loreti et al., 2008, Jeong et al., 2010, Qi et al., 2011, Das et al., 2012, Page et al., 2012, Li et al., 2014, Xie et al., 2016, Plumb et al., 2018, Wang et al., 2020, An et al., 2021). Although controversially discussed, ROS accumulation (during HL) positively or negatively impacts FB genes and anthocyanin accumulation. (Vandenabeele et al., 2004, Vanderauwera et al., 2005, Miller et al., 2007, Viola et al., 2016, Xu et al., 2017). FB genes are also stimulated by exogenous sugar supply (Creasy, 1968, Teng et al., 2005, Solfanelli et al., 2006), and starch-deficient mutants with high-sugar contents accumulate more anthocyanins than wild-type plants (Ragel et al., 2013, Schmitz et al., 2014). Despite no change in FB transcript accumulation was observed, knockout of *TRIOSEPHOSPHATE (TP)/PHOSPHATE TRANSLOCATOR* (*TPT*) essential for the immediate export of photoassimilates (Schneider et al., 2002), resulted in low anthocyanin level after HL treatment (Schmitz et al., 2014). Additionally, blocking the photosynthetic electron transfer by 3-(3,4-dichlorophenyl)-1,1-dimethyl urea (DCMU) suppresses FB (Schneider and Stimson, 1971, Akhtar et al., 2010, Jeong et al., 2010). These findings may directly connect photosynthetic activity and plastid-derived carbohydrates with FB regulation, particularly when light intensity increases. In this context, the analysis of plants with altered activity or content of central components of the sugar-signalling network suggests a function of these factors in FB regulation. For example, perturbation of the TARGET OF RAPAMYCIN (TOR) complex, which is involved in growth-promoting glucose-signalling (Xiong et al., 2013, Chen et al., 2018), results in high anthocyanin contents (Wang et al., 2017, Salem et al., 2018). Also, the function of a second major sugar-sensor SNF1-RELATED PROTEIN KINASE 1 (SnRK1, Baena-Gonzalez et al., 2007), which is activated under low energy-stress such as sugar starvation or hypoxia, is connected to FB. When cellular sugar levels are high, SnRK1 activity is repressed by regulatory sugar-phosphates, like trehalose-6 phosphate (T6P), glucose-1 phosphate or glucose-6 phosphate (Zhang et al., 2009, Nunes et al., 2013, Zhai et al., 2018). Repression of SnRK1 activity through increased production of T6P by overexpression of T6P SYNTHASE from *E. coli* (*otsA*) or by inducible knockdown of the second catalytic SnRK1 subunit (*KIN11*) in a *KIN10* knockout background activates FB even under normal growth conditions. In contrast, reduced anthocyanin contents were observed when T6P PHOSPHATASE (*otsB*) or the catalytic *KIN10* subunit was overexpressed. Intriguingly, the central FB TF *PAP1* is repressed when SnRK1 activity is high but induced after inactivation of SnRK1 (Schluepmann et al., 2003, Baena-Gonzalez et al., 2007, Zhang et al., 2009, Wingler et al., 2012, Nukarinen et al., 2016, Wang et al., 2021). SnRK1 also functions as a post-translational FB regulator by indirectly stabilizing PAL(s), the first enzyme in the cytosolic phenylpropanoid pathway (Wang et al., 2021).

Previous research revealed a complex network for the transcriptional regulation of FB depending on or interacting with various plastid-derived molecules and known components of other signalling pathways. However, the exact nature of the signal stimulating FB during HL acclimation is still unknown. Also, although TFs regulating *EBG* and *LBG* have been identified, molecular mechanisms and components perceiving, integrating, and translating the signal to changes in FB gene expression in response to increased light intensity are not well understood. Here we provide experimental evidence that photosynthetic activity, carbon fixation, immediate export of photoassimilates from chloroplast through TPT, and an increase of cellular sugar pools serve as a signal for HL induction of FB. Our results are in favor of SnRK1 as sensor and integrator of a sugar-signal downstream of chloroplasts regulating FB and likely other processes during HL acclimation.

## Results

### Perturbation of Triosephosphate export diminishes HL induction of FB

To analyze the effects of altered photoassimilate partitioning on the regulation of FB, we used *Arabidopsis thaliana* wild-type (WT, Col-0), starch-deficient *adg1-1* (Fig. S1A) and the *tpt-2* knockout mutant for the *TRIOSEPHOSPHATE (TP)/PHOSPHATE TRANSLOCATOR* (*TPT*) (Fig. S1B) in high light (HL) shift experiments. WT plants showed a 2-fold increase in glucose (GLC), fructose (FRC) and sucrose (SUC) contents after short-term HL treatment, whereas the deficiency of *TPT* prevented HL-induced accumulation of all three sugars (Fig. S1C-E). In agreement with previous studies (Schmitz et al., 2014), block of starch biosynthesis in *adg1-1* led to overaccumulation of GLC, FRC and SUC 8 h after HL shift relative to the WT. After HL exposure, *adg1-1* accumulated 2.5-fold more anthocyanins, but lack of *TPT* resulted in only 20% of the anthocyanin content compared to WT (Fig. 1B). Transcript analysis revealed a higher expression of the transcription factor (TF) *PAP1* and its regulated target involved in anthocyanin biosynthesis (*LDOX*, Fig. 1A) in *adg1-1* compared to WT after 8 h of HL treatment (Fig. 1C). Expression of *PAP1* and *LDOX* was significantly diminished in *tpt-2* compared to WT after 8 h of HL (Fig. 1E and F), but not after 24 h of HL (Fig. 1E and F). In contrast to *tpt-2*, sugar (phosphate) transporter mutants for GLUCOSE 6-PHOSPHATE/PHOSPHATE 2 (*gpt2-1*), XYLULOSE 5-PHOSPHATE/PHOSPHATE TRANSLOCATOR (*xpt-1 and 2*) and PLASTIDIC GLUCOSE TRANSLOCATOR (*pglct-2*) showed WT-like accumulation of anthocyanins after 24h HL (Fig. S1F). Collectively, these results indicate specific and dynamic changes in the expression of FB genes and, ultimately, the potential to accumulate anthocyanins during HL treatment, which was significantly reduced in *TPT* deficient mutants.

To analyze the induction of FB during HL in more detail, we performed a time-resolved analysis of anthocyanin, starch, sugar, and transcript abundances and harvested samples every three hours upon the HL shift of Col-0 and *tpt-2* (Fig. 2). Anthocyanin contents increased after 6-9 h HL in WT, however, they started to increase only after 15-18 h HL in *tpt-2* (Fig. 2A). Starch accumulation was similar between *tpt-2* and WT plants until 9-12h in the HL. After that time point, the *tpt-2* mutant continued to synthesize starch resulting in significantly higher levels at the end of the HL experiment (Fig. 2B). HL exposure resulted in a fast and robust accumulation of GLC and FRC within 3 h in WT leaves (Fig. 2C and D), but sugar contents only gradually increased in *tpt-2* and reached WT-like level after 12-15 h HL (Fig. 2C and D). The TF MYB111 and its targets, *CHS* and *FLS1*, were strongly induced in WT from the beginning of the HL shift. Although the trend was overall similar, *EBG* expression was diminished in *tpt-2* compared to WT (Fig. 2E-G). Intriguingly, HL induction of *LBG* involved in anthocyanins biosynthesis was significantly delayed in *tpt-2* by more than 8 h. While *PAP1* expression increased by 200-300 fold within 6 h of HL treatment in the WT compared to t(0), it remained low in *tpt-2* within the first hours of HL exposure and only started to increase after 9 h HL (Fig. 2H). Consequently, expression of *DFR* and *LDOX* was markedly stimulated between 3 h and 15 h and reached a plateau after 18 h HL incubation in WT plants (∼300-folds compared to t(0)). In *tpt-2*, the *LBG* expression increased to WT level only after 24 h of HL treatment (Fig. 2I and J). Noteworthy, the kinetic of *LBG* induction in *tpt-2* after 12 h of HL treatment resembled the induction in WT between 3-15 h HL. Hence, delayed stimulation of FB pathway genes agreed with the late and diminished anthocyanin accumulation (Fig. 1A and 2A) and paralleled the accumulation of sugars in *tpt-2*. In contrast to FB genes, *PHOTOSYNTHESIS ASSOCIATED NUCLEAR GENES* (*PhANGs*) were repressed similarly in WT and *tpt-2* upon HL exposure (Fig. 2K and L). Previously, ROS were proposed to function in FB regulation. However, no difference in ROS level between *tpt-2* and WT were observed using specific stains (Fig. S2C-E). We also found WT-like expression of stress- or ROS responsive genes, such as *GLUTATHIONE PEROXIDASE 7* (*GPX7*) or *ASCORBATE PEROXIDASE 1* (*APX1*), in *tpt-2* during HL (Fig. S2A and B).

**Figure 2:**
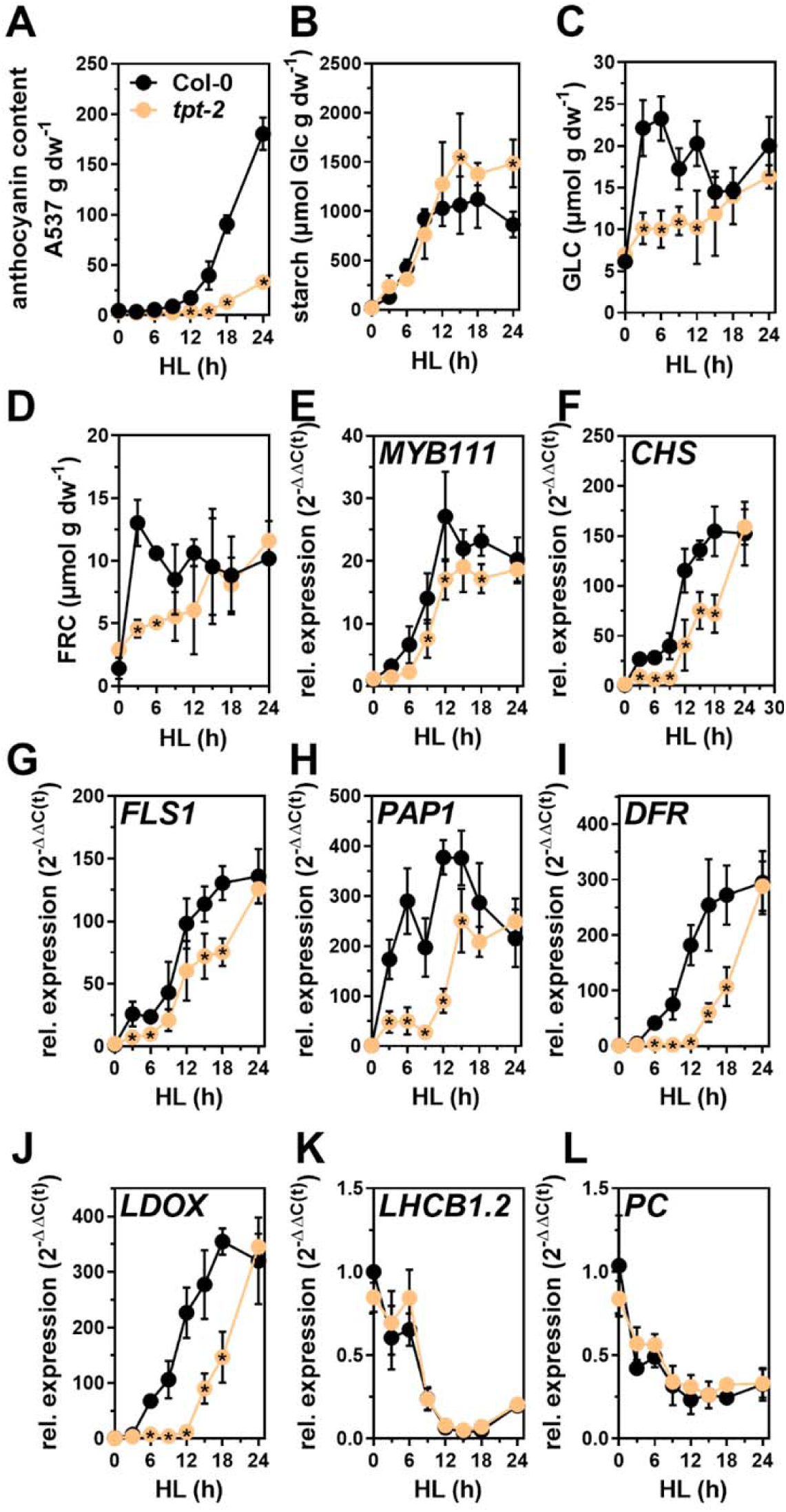
Kinetic analysis of (**A**) anthocyanin, (**B**) starch (represented as glucose equivalents), (**C**) glucose (GLC), (**D**) fructose (FRC) content and gene expression (**E-L**) in Col-0 (black) and *tpt-2* (orange) throughout a 24h HL shift experiment. Plants were shifted at the end of night (t0) to the HL condition, and samples were collected at the indicated time points. Gene expression was calculated using the 2^−ΔΔ(C(t))^ method relative to Col-0 (t0) and *SAND* as reference gene. Values are mean ±SD (n=4). Statistical significance between wild-type and *tpt-2* at each time-point was analyzed using student’s t-test (*p*<0.05). Significantly different values are indicated with asterisks inside the *tpt-2* symbols. (**E-G**) Expression of representative *EARLY BIOSYNTHETIC GENES* (*EBGs*), (**H-J**) *LATE BIOSYNTHETIC GENES* (*LBGs*) involved in FB and (**K-L**) *LIGHT HARVESTING CHLOROPHYLL A/B BINDING PROTEIN 1*.*2 (LHCB1*.*2)* and *PLASTOCYANIN (PC)* was analyzed.

Because *TPT* deficiency caused a substantial delay in the accumulation of sugars in the HL we hypothesized that the lack of a sugar signal originating from chloroplasts was causative for the repressed FB pathway genes in *tpt-2*. To test this, Col-0, *tpt-2* and *adg1-1* were incubated in buffer without or supplemented with sucrose and shifted to HL (Fig. 3). In the control buffer, *tpt-2* and *adg1-1* showed the expected deregulation of FB on the transcriptional and metabolic level as observed for soil-grown plants shifted to HL (Fig. 3A-C), but sucrose feeding fully restored the diminished expression of *PAP1* and *LDOX* (Fig. 3B and C) and anthocyanin contents (Fig. 3A) to WT-level in *tpt-2* in HL. Furthermore, overaccumulation of *PAP1* and *LDOX* transcripts in the sugar-accumulating *adg1-1* mutant (Fig. S1C-E) was suppressed by introducing the *tpt-2* allele after short term HL treatment (Fig. 3D and E). To provide additional evidence for TP biosynthesis in chloroplasts permitting FB activation, carbon fixation within the Calvin cycle was restricted by adjusting the CO_2_ concentration using a Li-Cor device. *PAP1* and *LDOX* were significantly induced after HL treatment at 400 ppm CO_2_ (Fig. 3F and G), and expression changes were comparable to those observed in young WT plants treated with bright LED light in a growth cabinet (for example, Fig. 2H for *PAP1*). In contrast, a limited CO_2_ supply of 50 ppm suppressed the HL induction of *PAP1* and abolished *LDOX* induction (Fig. 3F and G). We also found that plants shifted to 24h HL at different time points during short-day (SD) or long-day (LD) growth conditions accumulated more anthocyanins the later they were shifted during the day. These experiments revealed a positive correlation of FB with the starch content before the HL (Fig. S3A and B). Moreover, *PAP1* and *LDOX* expression after 8 h of HL and the anthocyanin contents after 24 h HL were also positively correlated with the sugar contents before the HL shift (t0) (Fig. S3 C-K). In other words, plants with higher sugar contents before the HL shift showed stronger FB gene induction and higher anthocyanin contents during HL. In summary, the kinetic analysis showed that proper induction of FB and accumulation of anthocyanins during HL acclimation depends on carbon fixation, TP export from chloroplasts and cellular sugar contents.

**Figure 3:**
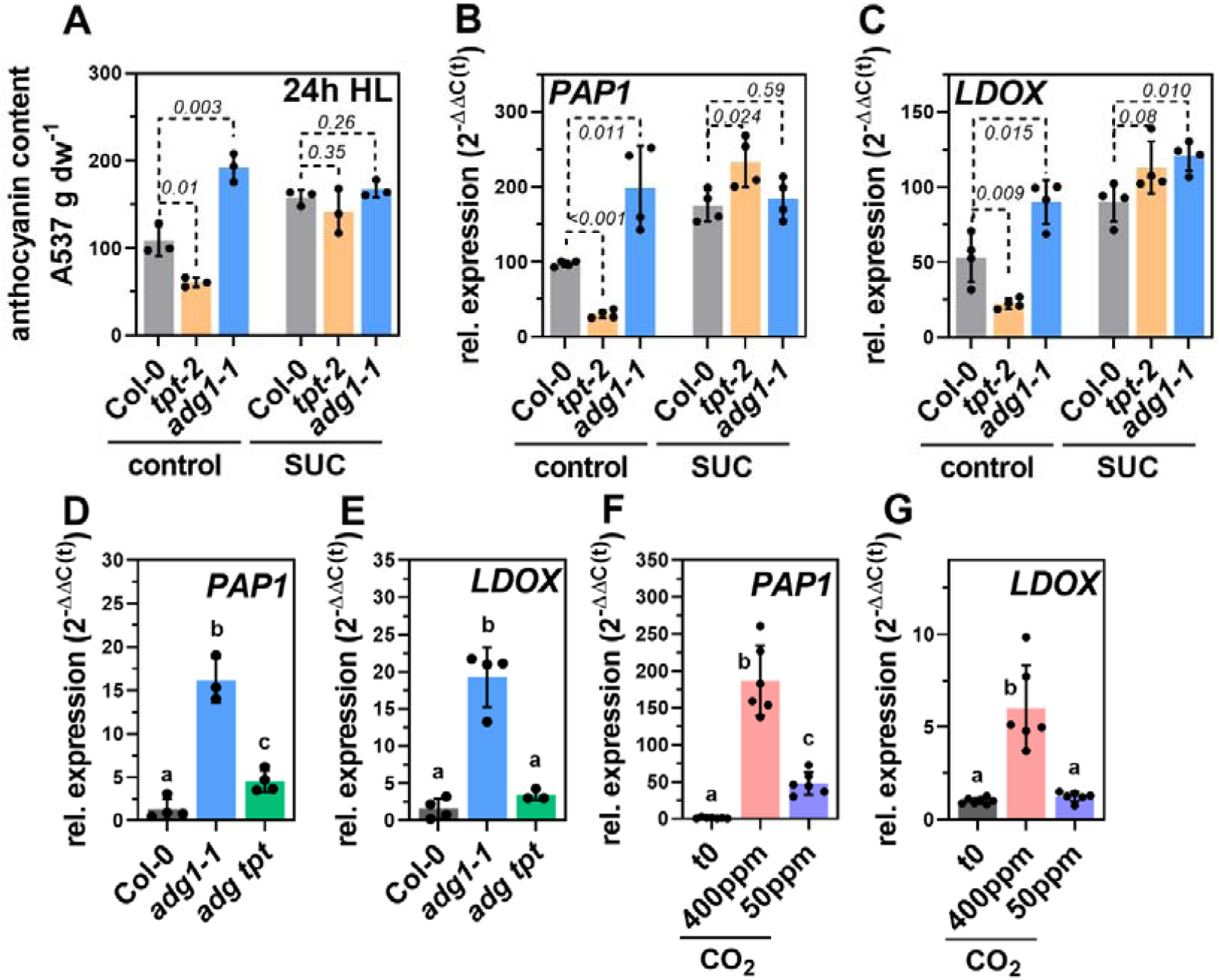
Induction of FB depends on carbon fixation, TP export and carbohydrates. (**A**) Anthocyanin accumulation and relative expression of (**B**) *PAP1* and (**C**) *LDOX* in Col-0 (grey), *tpt-2* (yellow) and *adg1-1* (blue) mutant in the absence (control) or presence of 5% sucrose (SUC). Seedlings were transferred to buffer +/-SUC at the EoN and exposed to HL for 8h (transcript analysis) and 24h HL (anthocyanin content). Gene expression was calculated using the 2^−ΔΔ(C(t))^ method relative to Col-0 before the transfer to the incubation buffer and *SAND* as reference gene. Values are mean ±SD. Statistical significance between wild-type and mutants in each condition was analyzed using student’s t-test, and the p-values are shown (n=4 for transcripts and n=3 for anthocyanin content). (**D**) Relative expression of *PAP1* and (**E**) *LDOX* in Col-0 (grey), *adg1-1* (blue) and *adg1-1 tpt-2* (green) after 8h HL treatment. Gene expression was calculated using the 2^−ΔΔ(C(t))^ method relative to Col-0 after 8h HL and *SAND* as reference gene. (**F**) Relative expression of *PAP1* and (**G**) *LDOX* in Col-0 before the incubation (EoN, t0, grey) in 400ppm CO_2_ (light pink) or 50ppm CO_2_ (purple). Leaves were mounted in a Li-Cor device and incubated for 4h at 500μmol photons m^−2^ s^−1^. Gene expression was calculated using the 2^−ΔΔ(C(t))^ method relative to the expression at t0 and *SAND* as reference gene. 5-week old rosette plants were used, and each data point represents one leaf sample from independent plants. Statistical significance between wild-type/control and mutants/treatment was calculated using one-way ANOVA (Dunnett’s multiple comparisons test) relative to Col-0 after 8h HL (in D and E, n≥3) or the expression values prior to the incubation at different CO_2_ concentrations (t0 in F and G, n=6). Values are mean ±SD.

### Transcriptome analysis revealed dynamic regulation of the HL acclimation response in *tpt-2*

To gain further insights into the HL acclimation response and its dependency on TP export from chloroplasts, we performed a whole transcriptome sequencing analysis (mRNA sequencing, mRNAseq) of WT and *tpt-2* after 9 h and 18 h of HL. We particularly selected these timepoints because the anthocyanin branch of FB was not induced in *tpt-2* compared to WT at 9 h of HL treatment. In contrast, after 18 h of HL, *LBG* expression reached a maximum in WT but was induced in *tpt-2* relative to 9 h HL (Fig. 2). Hence, ratiometric analysis of gene expression changes between these time points would also disclose other dynamic changes in the *tpt-2* transcriptome. From the initial comparison of WT and *tpt-2* (Fig. 4), we found only a minor overlap of commonly differentially expressed genes (DEG, adjusted p-value<0.05) between 9 h and 18 h HL (Fig. 4A, Fig. S4A and Table S2). While this already indicated a dynamic transcript expression in *tpt-2*, relative changes between mutant and WT were also influenced by changes in WT between 9 h and 18 h HL treatment (Fig. S4A). Gene Ontology Term Enrichment (GO term) analysis revealed “phenylpropanoid/flavonoid/anthocyanin biosynthesis and regulation”, “starch metabolic process”, and “cellular carbohydrate metabolic process” as most strongly down-regulated gene groups in *tpt-2* versus WT at 9 h HL (Table S2, log_2_ FC (fold change) <1). Among the down-regulated genes in *tpt-2* at 9 h HL, we found *GLUCOSE-6-PHOSPHATE/PHOSPHATE TRANSLOCATOR 2* (*GPT2*), whose expression was previously reported to depend on TP export from chloroplasts during HL (Weise et al., 2019). Furthermore, the large subunit of *ADP-GLUCOSE PYROPHOSPHORYLASE 3* (*APL3*) and *GRANULE BOUND STARCH SYNTHASE 1* (*GBSS1*), both involved in starch biosynthesis (regulation), were repressed in *tpt-2*. Noteworthy, components of light signalling pathways, such as *CRYs, PHYs* or components of the E3 ubiquitin ligase COP1/SPA, and downstream factors *HY5* and its interacting *BBX20-22*, were not repressed in *tpt-2* relative to WT throughout the HL kinetic (Table S1). Transcripts related to “trehalose metabolism” (*TREHALOSE-6-PHOSPHATE SYNTHASE, TPS8,9,10,11*) were upregulated in *tpt-2* at 9 h HL (log_2_ FC>1). Upregulated and down-regulated transcripts in *tpt-2*/WT at 9 h HL were also grouped to the “cellular response to oxygen levels/hypoxia” (Table S2).

**Figure 4:**
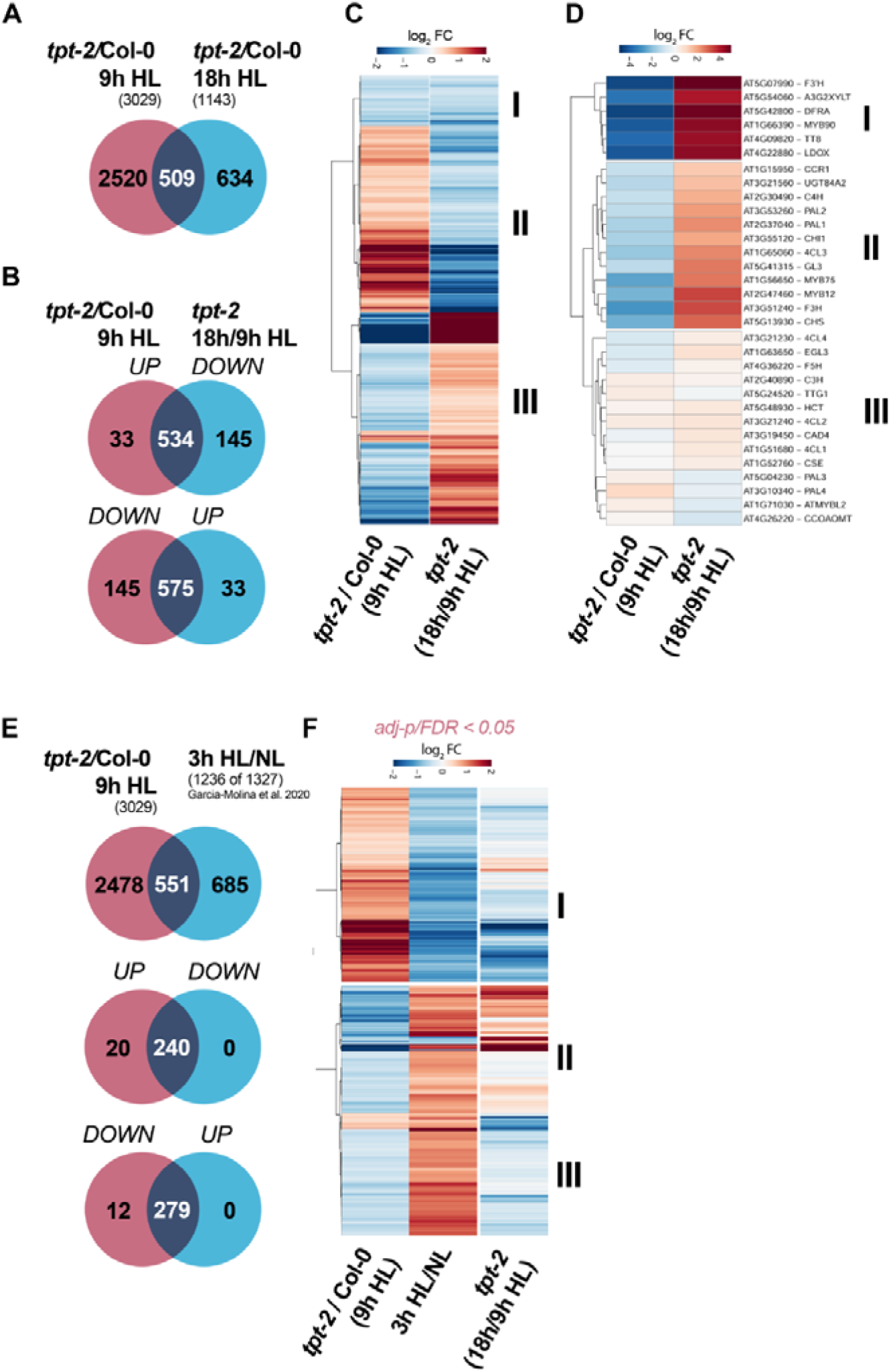
mRNA-seq analysis of HL-treated plants revealed a delayed HL response in *tpt-2*. **(A)** Venn diagram depicting the overlap of significantly differentially expressed genes (DEGs, adjusted p-value<0.05) between Col-0 and *tpt-2* incubated for 9 h (red) and 18 h (blue) in the HL. (**B**) Venn diagram and (**C**) heatmap showing transcripts differentially expressed in *tpt-2*/Col-0 at 9h HL and their change in expression after 18 h relative to 9 h HL in *tpt-2* (*tpt-2* 18h/9h HL) (adjusted p-value<0.05). A significant number of up-or downregulated DEGs in *tpt-2* relative to Col-0 after 9h HL were induced or repressed in *tpt-2* after 18 h of HL treatment (in total 1109). DEGs in Cluster I were downregulated in *tpt-2*/Col-0 after 9 h HL and were further repressed in *tpt-2* 18 h/9 h HL. Cluster II encompasses DEGs upregulated in *tpt-2*/Col-0 after 9h HL but significantly downregulated in *tpt-2* 18h/9h HL. Cluster III contains DEGs downregulated at 9h HL in *tpt-2* relative to Col-0 but were induced at 18h HL treatment in *tpt-2* relative to 9h HL. (**D**) Expression of genes encoding enzymes and transcriptional regulators of phenylpropanoid and flavonoid biosynthesis in *tpt-2*/Col-0 at 9h HL and relative changes in *tpt-2* at 18h compared to 9 h HL. Cluster I and II contained transcripts that were strongly suppressed in *tpt-2*/Col-0 at 9 h HL but were induced after 18h HL in *tpt-2* relative to 9h HL. Cluster III encompassed transcripts that showed only minor (relative) changes between 9h and 18h HL treatment of *tpt-2*. (**E** and **F**) Comparison of DEGs in *tpt-2*/Col-0 (9 h HL) with DEGs after a short term (3 h) HL shift of *Arabidopsis* WT plants. DEGs after 3h HL relative to normal light (NL) condition were extracted from Garcia-Molina et al. 2020 (false discovery rate (FDR) <0.05). Of the 1327 DEGs after 3h HL treatment, transcripts for 1236 genes were found in the *tpt-2* data set and used for comparison. (**F**) Heatmap comparing relative expression changes of transcripts in 3h HL treated Col-0 (Garcia-Molina et al. 2020), *tpt-2*/Col-0 (9 h HL) and *tpt-2* (18 h HL/9 h HL). Changes in gene expression are given as log_2_ fold change (FC) relative to the control and hierarchical row clustering (Euclidean distance) using ward.D method was applied for all heatmaps.

We focussed next on the dynamic changes in the *tpt-2* transcriptome by analyzing the list of DEG at 9 h HL for their expression change at 18 h HL (Fig. 4). Among the deregulated transcripts at 9 h HL, approximately 35% (1109 transcripts) showed opposite expression at 18 h HL in *tpt-2*, i.e. were induced or repressed (adjusted p<0.05). 534 DEG were upregulated in *tpt-2*/WT at 9 h HL but were repressed in *tpt-2* after 18 h relative to 9 h HL. 575 DEG repressed at 9 h HL relative to the WT were induced at 18 h HL in *tpt-2* (Fig. 4B and C, clusters II and III). Transcripts that we classified as late-induced in *tpt-2*, were assigned to GO terms related to “flavonoid biosynthesis” and “carbohydrate metabolic process” (Fig. 4D and Table S4, filtered for −1<log_2_ FC>1). In addition to *APL3, GBSS1*, and *GPT2* transcripts encoding FB enzymes, such as *DFR* and *LDOX*, and the TFs *MYB75* (*PAP1*), *MYB90* (*PAP2*), *MYB113, MYB114* and *TT8* were markedly induced after 18 h HL treatment in *tpt-2* (Fig. 4D cluster I and II, Table S5). On the contrary, *DARK INDUCIBLE 1* (*DIN1*/*SEN1/STR15*), *DIN10, TPS8* and *11*, regulated by SnRK1 signalling (Baena-Gonzalez et al., 2007, Zhang et al., 2009), were induced relative to WT at 9 h HL but late-repressed in *tpt-2* after 18 h HL.

In addition, we compared the *tpt-2* data set (9 h HL) with the result of previous transcriptome analysis of WT plants shifted to 3 h HL under similar conditions (Table S6, Garcia-Molina et al., 2020). We found a significant overlap of DEG between both sets (Fig. 4E and F). Of the commonly DEG, 50% were repressed in WT plants at 3 h HL but almost exclusively upregulated in *tpt-2*/WT after 9 h HL (Fig. 4E). Nearly all transcripts in this cluster were late-repressed in *tpt-2* at 18 h HL (Cluster I, Fig. 4F) and belonged to, for example, “carbohydrate metabolism” (including “trehalose metabolism”) and “response to hypoxia” (Table S6). On the contrary, 279 transcripts induced in WT plants after 3 h HL were downregulated at 9 h HL in *tpt-2* and part of them were late-induced in *tpt-2* after 18 h HL (Fig. 4F, Cluster II). These transcripts belonged to “carbohydrate metabolic process”, “protein folding”, and “ribosome biogenesis and function” (Table S6). In summary, in the absence of *TPT*, plants cannot induce transcriptome adjustments as observed for WT plants during early phases of HL acclimation (e.g., FB, enzymes of starch and carbohydrate metabolism or factors of protein translation) but, instead, switch to the HL state only after prolonged HL treatment.

It has been previously shown that HL acclimation also involves and depends on the transcriptional adjustment of phytohormone biosynthesis (Huang et al., 2019), and phytohormones are involved in the transcriptional and post-translational regulation of FB. Hence, we analyzed the RNAseq data set for the expression of gene products involved in phytohormone biosynthesis and signalling (Table S3, −1>log_2_ FC>1, adj-p<0.05). From the group of auxin-responsive genes analyzed, only *SMALL AUXIN-UP RNA* (*SAUR*) *21,20,22* and *64* were moderately upregulated in *tpt-2* throughout the HL treatment. Among the ABA-related transcripts tested, *PYL5* and *PYL7* were moderately induced at 9 h HL in *tpt-2*/WT (2-fold), and *PYL7* was repressed after 18 h HL in *tpt-2* (Table S3). Two transcripts related to jasmonate (JA) biosynthesis were deregulated in *tpt-2*/WT at 9 h HL. *ACYL-COA OXIDASE 4 (ACX4)* was significantly upregulated, and *JASMONIC ACID CARBOXYL METHYLTRANSFERASE (JMT)* was downregulated, and both transcripts showed opposite expression in *tpt-2*/WT relative to values published for an HL acclimation kinetics of WT plants (Fig S4B). Additionally, we found that JA biosynthesis genes, particularly gene products for precursor biosynthesis inside chloroplast (*LIPOXYGENASE, ALLENE OXIDE SYNTHASE* and *ALLENE OXIDE CYCLASE*), were strongly and specifically induced in *tpt-2* at 18 h HL relative to 9 h HL (Fig. S4C, Table S3). Although JA promotes anthocyanin biosynthesis and FB pathway was induced in *tpt-2* after long-term HL, the JA-deficient *jassy* mutant (Guan et al., 2019) showed WT-like anthocyanin accumulation during HL treatment (Fig. S4D). These analyses revealed that hormone-related gene expression changes in *tpt-2* after long-term HL but likely plays a less important role for the observed changes in FB kinetics compared to sugar signalling.

### The impact of SnRK1 on HL induction of FB

The gene expression analyses identified a large set of genes that were strongly upregulated (9 h HL) but late-repressed (18 h HL) in *tpt-2*, which were reported to be regulated by SnRK1 (Fig. 5, Table S7). The comparison with previously conducted transcriptome analysis (Baena-Gonzalez et al., 2007) revealed that 50% of the transcripts deregulated by *KIN10* overexpression in *Arabidopsis* protoplasts (*KIN10ox*) were also deregulated in *tpt-2* after 9 h HL treatment (Fig. 5A and Table S7). Intriguingly, these transcripts were almost exclusively deregulated in the same direction in *KIN10ox* and *tpt-2* after 9 h HL but were overall repressed or induced, respectively, after 18 h HL in *tpt-2* (Fig. 5B). Furthermore, the *tpt-2* transcriptome at 9 h HL showed a positive correlation with changes in gene expression induced by *KIN10ox* and starvation conditions (Fig. 5B and S5A). After prolonged HL treatment, expression changes in *tpt-2* were positively correlated with those observed after sucrose and glucose feeding (Fig. 5B and S5A). The late-repressed DEG in *tpt-2* were related to GO terms for “carbohydrate metabolism”, “catabolic process”, “starvation”, and “absence of light” (Table S7), and late-induced transcripts were mainly grouped to “ribosome assembly and function” and “protein translation” (Table S7). In summary, the analysis revealed dynamic changes in the expression of transcripts responsive to carbohydrate availability and pointed to SnRK1 as a potential downstream component of sugar signalling during HL treatment.

**Figure 5:**
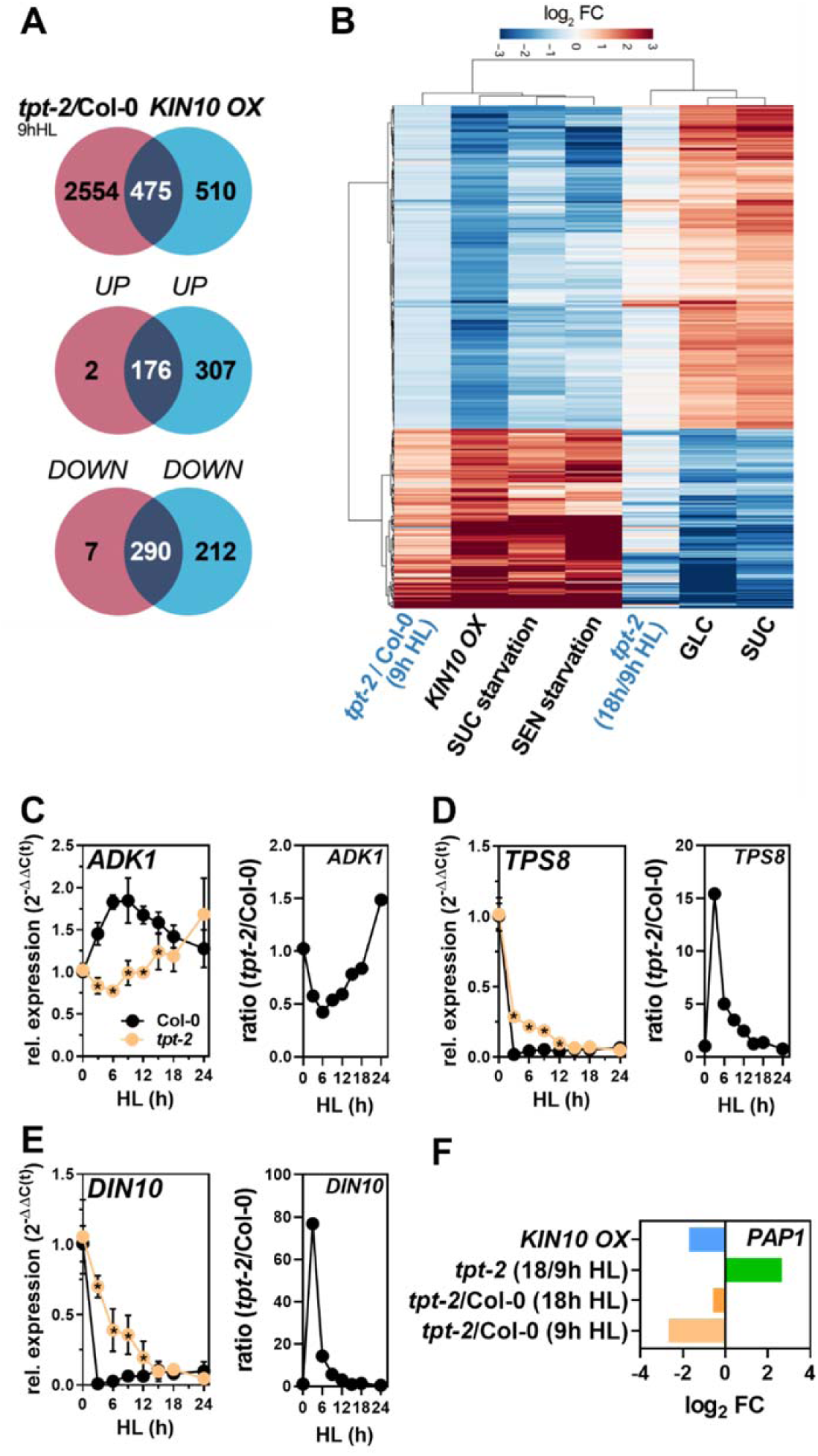
Comparative transcriptome analysis indicated deregulated SnRK1 signalling in *tpt-2* during HL acclimation. (**A**) Venn diagram showing the overlap between DEGs in *tpt-2*/Col-0 after 9h HL and overexpression of *KIN10* in protoplasts (*KIN10 OX*) extracted from Baena-Gonzales et al. 2007 (Table S7). Of the 985 reported transcripts for *KIN10 OX*, 475 were deregulated in the same direction in *tpt-2* after 9 h HL (compare Venn diagrams for UP/UP and DOWN/DOWN). (**B**) Heatmap comparing the relative expression of DEGs in *tpt-2*/Col-0 9 h HL (adjusted p-value<0.05) and in *tpt-2* (18 h/9 h HL) with the expression changes induced by *KIN10* overexpression in protoplasts, sucrose (SUC) and senescence (SEN) starvation, glucose (GLC) and sucrose (SUC) feeding (Table S7). Changes in gene expression are given as log_2_ fold change (FC) relative to the control in each data set. Hierarchical column and row clustering (Euclidean distance) using ward.D method were applied. (**C-E**) Relative expression of SnRK1-repressed (C) *ADENYLATE KINASE1*, and SnRK1-induced (D) *TREHALOSE-6-PHOSPHATE SYNTHASE 8* and (E) *DARK INDUCIBLE 10/ RAFFINOSE SYNTHASE* (*DIN10*) during the HL shift kinetic of Col-0 and *tpt-2*. Gene expression was calculated using the 2^−ΔΔ(C(t))^ method relative to Col-0 (t0) and *SAND* as reference gene. Statistical significance between wild-type and *tpt-2* at each time-point was analyzed using student’s t-test (n=4, *p*<0.05, asterisk inside the *tpt-2* symbols). Values are mean ±SD. (**F**) Expression of *PAP1* (encoded by *MYB75*) in the indicated data sets/comparisons. Expression values for *PAP1* in protoplasts overexpressing *KIN10* were extracted from previously published data (Baena-Gonzales et al. 2007). Changes in gene expression are given as log_2_ fold change (FC) relative to the control in each data set. *PAP1* was significantly deregulated in *KIN10 OX* relative to control protoplasts, *tpt-2*/WT (9h HL) and *tpt-2* 18h/9h HL.

To further test if SnRK1-dependent regulation of gene expression was compromised in *tpt-2* during HL treatment, we performed a targeted analysis of transcripts deregulated by *KIN10ox* using the sample set of the HL kinetic (Fig. 2). *ADENOSINE KINASE 1 (ADK1)*, repressed by *KIN10ox*, was significantly induced in WT plants within the first hours but was not induced in *tpt-2* until 12 h HL treatment (Fig. 5C). In contrast, two SnRK1 induced transcripts, *TPS8* and *DIN10*, were repressed within 3 h HL treatment of WT plants, but repression was markedly delayed in *tpt-2* (Fig. 5D-E). The catalytic SnRK1 subunit *KIN10* and other subunits in *tpt-2* were not differentially expressed between WT and *tpt-2* (Fig. S5B, Table S1), pointing to a post-translational mechanism leading to high SnRK1 (early time points) and repressed SnRK1 (late time points) activity, respectively. Because overexpression of the catalytic SnRK1 subunit *KIN10* represses *PAP1* in protoplasts assays (Baena-Gonzalez et al., 2007) and *PAP1* showed dynamic changes in expression in *tpt-2* throughout the HL treatment (Fig. 5F and Fig. 2H), we hypothesized that SnRK1 is also involved in the regulation of FB during HL treatment (Fig. 6). SnRK1 activity is repressed by regulatory sugar-phosphates such as T6P. Overexpression of a *T6P PHOSPHATASE* (encoded by *otsB*, Fig. S6B) led to diminished expression of *PAP1* and *LDOX* during early time points of HL treatment (Fig. 6C-D) and significantly less anthocyanins compared to WT seedlings after 24 h HL (Fig. 6E). Induction of *PAP1* and *LDOX* was repressed in two independent *KIN10ox* lines compared to Ler WT plants (Fig. 6A-B, Fig. S6A), indicating that excess of KIN10 repressed FB genes during short term HL exposure. Consequently, *KIN10ox* lines accumulated only approximately 30% of the anthocyanins observed for the WT after 24 h HL (Fig. 6E). To test for the effect of SnRK1 suppression, an inducible RNAi line for *KIN11* (*SnRK1α2*), the second catalytic subunit of SnRK1, in the background of a *KIN10* (*SnRK1α1*) knockout mutant (*snrk1α1-3* amiRNAi *KIN11*, Pedrotti et al., 2018) was analyzed. This line had to be used because constitutive *snrk1α1/α2* are not viable, and single knockout results in no obvious phenotype (Baena-Gonzalez et al., 2007, Nukarinen et al., 2016, Belda-Palazon et al., 2020). After spraying ß-estradiol (ß-EST) using previously established protocols (Nukarinen et al., 2016), KIN11 was efficiently suppressed (Fig. S6C) and SnRK1 induced *DIN10* was downregulated in the ß-EST-treated plants compared to control plants at the end of the night (Fig. S6D). After subjecting to HL, +ß-EST plants had a visibly darker appearance, which was attributed to higher anthocyanin contents (Fig. 6F-G) and faster and more pronounced induction of *PAP1* and *LDOX* expression compared to control plants after the HL shift (Fig. 6H-I).

**Figure 6:**
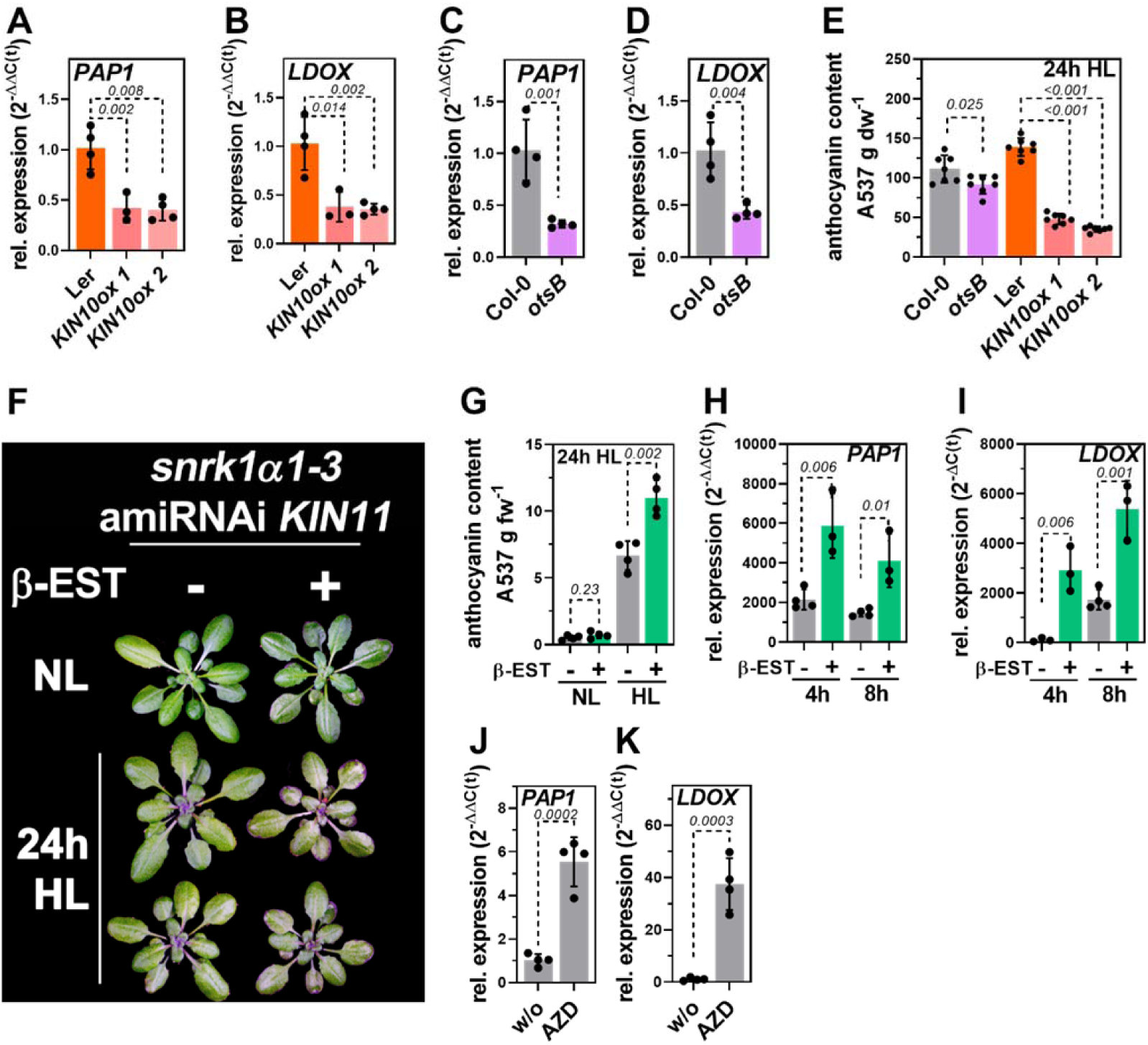
Expression of (**A**,**C**) *PAP1*, (**B**,**D**) *LDOX* in Ler (orange) and two independent overexpression lines for *KIN10* in Ler background (ox1 and ox2, pink and light pink) and Col-0 and Col-0 expressing *otsB* (light purple) after 8 h HL. Plants were shifted at the end of the night. Gene expression was calculated relative to the corresponding WT background after 8h HL using the 2^−ΔΔ(C(t))^ method and *SAND* as reference gene. Values are mean ±SD (n≥3), and *p-values* for student’s t-test between mutant and WT background are provided. (**E**) Anthocyanin accumulation in the lines shown in (A-D) after 24 h HL determined from two independent HL shift experiments. Values are mean ±SD (n=7) and p-values for student’s t-test between mutant and the corresponding WT background are shown. Statistical significance was analyzed using student’s t-test (n=4) and p-values are shown. (**F**) Phenotype and (**G**) anthocyanin content after 24 h HL, (**H**) *PAP1* and (**I**) *LDOX* expression in *snrk1α1-3* amiRNAi *KIN11* mutant where the knockdown of *KIN11* was induced by ß-estradiol (ß-EST) application. After three weeks of growth in SD, plants were sprayed without (−) or with 20 μM ß-EST (+) for another 7 days. Plants were shifted to the HL at the end of the night after the last ß-EST application. Gene expression was calculated relative to −ß-EST at EoN using the 2^−ΔΔ(C(t))^ method and *SAND* as reference gene. Statistical significance was analyzed using student’s t-test (n≥3) and p-values are shown. (**J**) Relative expression of *PAP1* and (**K**) *LDOX* in Col-0 after 6h HL treatment in the absence (w/o) or presence of 1 μM AZD8055 (AZD). Seedlings were grown on MS plates for 10d in SD conditions and shifted to HL at the end of the night. Gene expression was calculated relative to the control (w/o) using the 2^−ΔΔ(C(t))^ method and *SAND* as reference gene. Statistical significance between control and AZD treatment was analyzed using student’s t-test. Values are mean ±SD (n=4) and the *p*-values are shown.

In contrast to mutants with altered SnRK1 activity or content, the *HXK1* mutant *gin2-1* defective in GLC-mediated regulation of gene expression (Moore et al., 2003) accumulated WT-like amounts of anthocyanins in the HL (Fig. S6E). Inactivation of SnRK1 during HL shifts or increased sugar contents may also activate TOR (Nukarinen et al., 2016), affecting FB downstream of a sugar signal during HL treatment. To test the involvement of TOR, we grew Col-0 seedlings in the absence or presence of the TOR inhibitor AZD8055 and shifted them to HL. Intriguingly, inhibition of TOR further increased the expression of *PAP1* and *LDOX* in HL treated Col-0 (Fig. 6J-K). MAP kinases 3 and 6 phosphorylation depends on SnRK1 (Cho et al., 2016), and MPK3, 4, 6, and 11 have been shown to phosphorylate PAP1 (Li et al., 2016, Kreynes et al., 2020). Furthermore, MPK6 functions in the immediate regulation of gene expression downstream of TP export upon HL shifts (Vogel et al., 2014). However, two allelic *mpk6* mutants were not affected in anthocyanin accumulation during a 24 h HL shift (Fig. S6F).

## Discussion

### Carbon fixation, TP export and increase of cellular sugar level are essential for HL induction of FB

The time-resolved targeted and global gene expression analysis of HL treated plants disclosed a dynamic regulation of the HL-induced transcriptome response in *tpt-2*. Based on the presented results, we propose that *TPT* deficiency did not render the plants entirely insensitive to the HL, but rather dynamic changes in the signal connected to TP export from chloroplasts caused delayed stimulation of FB genes in *tpt-2*. Several lines of evidence support a role for TP export from chloroplasts as a metabolic retrograde signal and cellular sugar contents in the induction of FB pathway genes and accumulation of anthocyanins during HL. First, in WT seedlings, stimulation of FB genes and anthocyanin accumulation during HL treatment was positively correlated with cellular sugar contents before the HL shift, and limitation of carbon fixation significantly attenuated the activation of FB during short-term HL exposure (Fig. 3). In agreement with this, mutants impaired in photosynthesis due to perturbed Chl accumulation showed lower amounts of anthocyanin after HL compared to WT (Richter et al., 2020). Also, perturbation of photosynthetic activity by 3-(3,4-dichlorophenyl)-1,1-dimethylurea (DCMU) led to diminished expression of FB genes and anthocyanin contents in various species, which can be at least partially rescued by sugar feeding (Akhtar et al., 2010, Jeong et al., 2010, Su et al., 2016). Second, genetic perturbation of starch biosynthesis resulted in pronounced accumulation of soluble sugars (Fig. S1C-E), stimulated expression of FB pathway genes and anthocyanin production in HL-treated *adg1-1* (Fig. 1 and 3, Blasing et al., 2005, Ragel et al., 2013, Schmitz et al., 2014). Limited TP export in *adg1-1 tpt-2* double mutant prevented the rise in cellular sugar level (Schmitz et al., 2014) and diminished the induction of FB pathway genes after short-term HL treatment (Fig. 3). Lastly, the *tpt-2* mutant with reduced sugar contents after short-term HL (Fig. S1 and Fig. 2) failed to rapidly activate FB and accumulated only low anthocyanin contents but was fully rescued by sucrose feeding in HL. WT-like starch accumulation (Fig. 2), photosynthetic activity (Schmitz et al., 2014) and ROS-response (Fig. S2) of HL-treated *tpt-2* excluded overall perturbed photosynthesis and carbon fixation as a reason for the deregulation of FB in the mutant. Instead, these findings revealed sugar deficiency as causative for the lack of rapid FB induction in *tpt-*2 in response to HL. Consequently, a function for TP export from chloroplasts and subsequent conversion in cytosolic sugar biosynthesis pathways to activate FB genes during HL is strongly supported. This assumption is in good agreement with the PAP1-dependent positive impact of sugars on the expression of FB genes even in standard growth conditions (Teng et al., 2005, Solfanelli et al., 2006). Transcripts induced under energy-limiting conditions were enriched among the DEG during early time points of the HL shift, and sugar-responsive transcripts were late-responsive in *tpt-2* (Fig. 5 and S5). For example, *GPT2* was markedly repressed at 9 h HL relative to WT but strongly upregulated after 18 h HL in *tpt-2. GPT2* is induced in GLC-fed potato leaves (Quick et al., 1995) or by high sugar contents in starch biosynthesis mutants *adg1* and *pgm* (Kunz et al., 2010, Weise et al., 2019) and upon HL treatment of Arabidopsis WT plants (Huang et al., 2019). Hence, also the late induction of FB genes is most likely explained by the late increase of sugar contents in *tpt-2* during HL (Fig. 2), thereby supporting the suggested model of sugar-dependent regulation of FB during HL.

Considering the lack of the primary route for TP export, the question arises through which alternative pathway/transporter cytosolic sugar pools can increase in HL treated *tpt-2*. WT-like anthocyanin accumulation in *gpt2, pglct* and *xpt* (Fig. S1F) suggests that these routes for carbohydrate exchange are not essential for HL induction of FB in the presence of a functional TPT. Nevertheless, we cannot exclude that one of these transporters partially compensated for the lack of TPT in *tpt-2*. For example, XPT was reported to also transport TP (Eicks et al., 2002) and could (partially) complement the lack of TPT in HL as it does under normal growth conditions in *tpt-2* or *adg1-1 tpt-2* double mutants (Hilgers et al., 2018). Alternatively, starch degradation already occurring in *TPT*-deficient mutants during the day (Hausler et al., 1998, Schneider et al., 2002, Walters et al., 2004), and subsequent export of maltose through MEX1 (Niittyla et al., 2004) and conversion in the cytosol could also contribute to the rising sugar contents in *tpt-2* exposed to long-term HL. Indeed, a function of MEX1 for cold-induced anthocyanin accumulation was reported (Purdy et al., 2013), and maltose feeding can stimulate FB (Teng et al., 2005). Future analyses of double mutants deficient of *TPT* and other plastid-localized sugar transporters could reveal which alternative export pathways are responsible for the late increase in sugar contents and sugar-dependent regulation of FB in long-term HL.

Previously, ROS were shown to negatively or positively affect the expression of FB genes through, for example, redox-dependent regulation of transcriptional FB repressors such as TCP14 or 15 (Viola et al., 2016). However, the time-resolved analysis of WT and *tpt-2* mutant allowed the separation of the (early) ROS response from a metabolic sugar-signal regulating FB during HL acclimation. More precisely, expression of *GPX7* was rapidly stimulated upon HL exposure (3 h HL), declined and strongly increased again after 9 h HL in WT and *tpt-2* (Fig. S2). While FB gene expression parallels that of ROS marker genes during early time points in WT, *tpt-2* showed a WT-like ROS response but, at the same time, failed to stimulate crucial positive regulators (*PAP1, TT8*) of *LBG* (*LDOX, DFR*). The substantial increase of *GPX7* (and others, Fig. S2) after prolonged HL paralleled the expression of FB genes in *tpt-2* but not in WT, where the expression of FB pathway genes reached a maximum or was even repressed in the case of *PAP1*. In this connection, unaltered expression of ascorbate biosynthesis genes (e.g., *VTC2*, Table S1) and WT-like total ascorbate level in the *tpt-2* mutant (Schmitz et al., 2014) may exclude altered ascorbate level as causative for the deregulation of FB in *tpt-2* and, in turn, for the fast HL induction of FB in WT in our conditions (Page et al., 2012, Plumb et al., 2018). We do not rule out a function of ROS in the regulation of *EBG* (e.g., *CHS, FLS1*) or *LBG* in long-term HL (e.g., several days), under more severe or other abiotic stress conditions, but we concluded that ROS are most likely not involved in the stimulation of, particularly, *LBG* at early timepoints of HL treatment. In addition to ROS, phytohormones such as JA (Shan et al., 2009, Qi et al., 2011) or ethylene (Jeong et al., 2010) impact FB. Ethylene biosynthesis or responsive genes and other phytohormone biosynthesis pathways were not enriched among the DEG, and only JA biosynthesis genes were late-induced in HL treated *tpt-2* (Fig. S4B and C, Table S3). However, the JA-deficient *jassy* mutant (Guan et al., 2019) accumulated WT-like amounts of anthocyanins in HL (Fig. S4D), indicating that JA availability does not determine rapid activation of FB in HL.

At this point, we want to highlight that diminished anthocyanin accumulation in *tpt-2* during HL exposure had been previously reported by Schmitz et al. (2014). However, using microarray-based transcriptomics, the authors could not correlate changes in anthocyanin contents with changes in the expression of FB pathway genes. While we cannot entirely exclude that the different experimental conditions (shift to 300 μmol m^−2^ s^−1^ and another light source) had an impact on the different outcome of the experiments, the time point of harvest might explain most of the differences: Schmitz et al. harvested plant material after 4h and 48 h of the HL shift. However, our time-resolved analysis revealed only minor differences in the expression of FB pathway genes between WT and *tpt-2* after 4 h HL and (again) WT-like expression of *PAP1, DFR* and *LDOX* after prolonged HL exposure (i.e., 24 h HL) (Fig. 1 and 2). Hence, due to the selection of time points, the previous study could not detect the dynamic changes and altered transcriptional response during HL acclimation in *tpt-2*.

### Evidence for SnRK1 functioning downstream of chloroplasts for the induction of FB during HL acclimation

Transcriptome changes throughout the HL revealed late-induction and –repression of sugar-responsive genes in *tpt-2*, and we propose that increasing cellular sugar contents serve to rapidly induce FB on the transcriptional level upon HL exposure of plants. Changes in sugar contents are sensed and translated into adjustments of gene expression by, for instance, HXK1, TOR and SnRK1 and downstream components (Rolland et al., 2006, Baena-Gonzalez and Lunn, 2020). Our analysis showed that HXK1-dependent GLC-signaling is not essential for FB activation when plants are exposed to HL (Fig. S6E). Instead, the results favor a function of SnRK1 in HL induction of FB. During HL, overexpression of *KIN10* and catabolism of the SnRK1 inhibitor T6P in *otsB* (*T6P PHOSPHATASE)* diminished the induction of FB genes and the potential to accumulate anthocyanins (Fig. 6). On the other hand, induced knockdown of SnRK1 resulted in stronger and faster activation of pathway genes and anthocyanin accumulation (Fig. 6). Consequently, SnRK1 must be inactivated to allow the stimulation of FB during HL. This conclusion is supported by the targeted and global gene expression analysis (Fig. 4 and 5), showing altered SnRK1-dependent signalling in *tpt-2* in the course of the HL experiment. Based on the expression profile of well-established marker genes for SnRK1-dependent signaling, it can be concluded that the delayed increase of sugar contents (Fig. 2) led to the delayed inactivation of SnRK1 thereby permitting the activation of FB only after prolonged HL treatment of *tpt-2*. Likewise, when exogenous feeding artificially increased sugar levels (Fig. 3), fast induction of FB could be restored in *tpt-2*. In this regard, more pronounced activation of FB in a*dg1-1* during HL (Fig. 1 and 3) is likely explained by high sugar (Fig. S1) and T6P contents in starch-deficient mutants (Carillo et al., 2013), leading to (stronger) suppression of SnRK1 activity compared to WT plants. The newly established connection between FB activation in HL and SnRK1 activity gets strong support from previous studies: *PAP1* is repressed by overexpressing *KIN10* in protoplast assays and excess of *KIN10* diminished sucrose-induced activation of FB *in planta* (Baena-Gonzalez et al., 2007). On the contrary, FB is activated, and pathway genes are de-repressed in partial loss-of-function *snrk1α1*^−*/*−^*snrk1α2*^*+/*−^ and *snrk1α1-3* amiRNAi *KIN11* mutants even under normal growth conditions (Nukarinen et al., 2016, Peixoto et al., 2021, Wang et al., 2021). Overexpression of *T6P PHOSPHATASE* leading to the activation of SnRK1 in stably transformed lines (*otsB*, Zhang et al., 2009) repressed, but increased T6P level in *T6P SYNTHASE* (*otsA*) overexpressor plants induced FB (Schluepmann et al., 2003, Zhang et al., 2009, Wingler et al., 2012). In addition to the transcriptional regulation of FB pathway genes, the inactivation of SnRK1 leads to indirect post-translational stabilization of PHENYLALANINE AMMONIA-LYASE (PAL), which catalyzes the initial step of phenylpropanoid biosynthesis upstream of FB (Wang et al., 2021). Based on the new results presented, we propose that anthocyanin accumulation during HL acclimation is the result of both transcriptional and post-translational activation of FB through SnRK1 inhibition resulting from rapidly increasing sugar contents in HL.

TOR activity is stimulated by sugars (Xiong et al., 2013, Li and Sheen, 2016) and repressed by SnRK1-dependent phosphorylation of the RAPTOR component (Nukarinen et al., 2016). Therefore, high sugar contents and inactivation of SnRK1 during HL could result in the activation of TOR which may induce FB. However, diminished TOR activity in *raptor1B* or inducible *tor* knockdown mutants results in increased flavonoid contents (Caldana et al., 2013, Wang et al., 2017, Salem et al., 2018) and the application of the TOR inhibitor AZD even further stimulated rather than repressed FB pathway genes in HL (Fig. 6). While this suggests that active TOR is not essential for HL stimulation of FB, it may also point to a repressive function of TOR on FB (during HL treatment). On the other hand, perturbed TOR function leads to starch and sugar accumulation (Caldana et al., 2013, Salem et al., 2018, Zhang et al., 2018). Due to the strong correlation between sugar, mainly sucrose, and T6P contents (Carillo et al., 2013, Yadav et al., 2014, Peixoto et al., 2021), it could be hypothesized that FB is (further) stimulated when TOR activity is suppressed (in HL) because high sugar levels may result in stronger inactivation of SnRK1 (like in the sugar over accumulating starch mutants). The involvement of TOR and its interconnection with SnRK1 in the regulation of FB during HL acclimation should be analyzed in the future using (double) mutants with altered activities of both factors.

### Potential downstream components of the sugar signal and the connection with light-signaling

MPK6 was shown to be activated by a TP-dependent signal, thereby permitting the adjustment of nuclear gene expression within 10 min of HL exposure (Vogel et al., 2014). Although still controversially debated, phosphorylation of PAP1 by MPK3, 4, 6, or 11 may adjust the stability or activity of the TF and eventually the accumulation of anthocyanins (Li et al., 2016, Kreynes et al., 2020, Kreynes et al., 2021, Yang et al., 2021). Interestingly, phosphorylation of MPK3 and 6 depends on SnRK1 in response to submergence of leaves (Cho et al., 2016), which would provide a connection between TP-export, SnRK1, MPK activity, and the regulation of FB. According to our new results, however, SnRK1 needs to be inactivated to induce FB, and consequently, MPK6 would be not phosphorylated. In this context, *mpk6* mutants were indistinguishable from WT plants in terms of anthocyanin accumulation (Fig. S6F and Li et al., 2016), indicating that at least MPK6 is dispensable for HL-induced anthocyanin accumulation.

Compelling evidence for FB regulation by components of light-signalling pathways has been provided in the past. Thus, the question arises how SnRK1 may interact with light-signalling pathways to regulate FB during HL. CRY- and PHY-dependent inactivation of the CONSTITUTIVE PHOTOMORPHOGENIC1/ SUPPRESSOR OF PHYA-105 (COP1/SPA) containing E3 ubiquitin ligase complex in the light permits the stabilization of HY5 and its interacting B-box containing proteins (BBX20, BBX21 and BBX22, Bursch et al., 2020) but also PAP1 and PAP2 (Maier et al., 2013, Ponnu et al., 2019) and direct binding of these TFs to their target genes promote FB (Shin et al., 2013, Bursch et al., 2020). Consequently, *cry1, hy5* and *bbx202122* mutants showed low expression of FB regulators and pathway genes and diminished anthocyanin contents when exposed to monochromatic but also high light (Kleine et al., 2007, Stracke et al., 2010, Ponnu et al., 2019), thereby resembling mutants with active SnRK1 in terms of FB regulation (see above). Because physical interaction of SnRK1 with components of the cellular machinery for ubiquitin-mediated protein turnover is reported (E3 ligases and 26S proteasome, Farras et al., 2001, Nietzsche et al., 2014), it is tempting to speculate that SnRK1 directly or indirectly represses positive regulators of FB (such as PAP1, PAP2, MYB113, MYB114 or TT8). The positive regulator HY5 was recently reported to be phosphorylated by SPA proteins (Wang et al., 2021), but SnRK1 may also regulate the stability or activity, respectively, of HY5, the interacting BBX proteins or other TFs upstream of the transcriptional FB regulators and pathway genes. Despite not being differentially expressed in our analysis, NAC TFs acting as inducers (ANAC078, Morishita et al., 2009) or repressors (ANAC042 and ANAC032, Mahmood et al., 2016) of FB could be controlled by SnRK1. Although direct phosphorylation could not be demonstrated yet, the leucine zipper TF STOREKEEPER RELATED 1 (STRK1) is stabilized by SnRK1 overexpression in transient transformation assays, and *PAP1* expression and anthocyanin accumulation was repressed during late growth stages in *STRK1* overexpressor lines (Nietzsche et al., 2018). Hence, STRK1 would be a promising candidate for the regulation of FB during HL, acting downstream of SnRK1 and upstream of FB genes. A potential function of SnRK1 as a repressor of the highly energy-demanding FB is supported by, for example, SnRK1-mediated de-stabilization of TFs for fatty acid biosynthesis (Zhai et al., 2018) or thermoresponsive hypocotyl elongation (Hwang et al., 2019). Future experiments will reveal how and at which point light- and sugar-dependent signalling pathways converge on the molecular level to permit the activation of FB but also other (sugar-regulated) pathways through SnRK1 during plant acclimation to adverse environmental conditions.

Taken together, and in conjunction with previous studies disclosing a crucial function of TP export for the regulation of nuclear gene expression in (short-term) HL (Vogel et al., 2014, Weise et al., 2019), our study emphasizes the important role of chloroplasts as sensors for changes in the environment and emitter of metabolic signals for the adjustment of nuclear gene expression and pathways relevant for acclimation responses - such as FB. Induction of FB and accumulation of anthocyanins is a widespread response of plants exposed to adverse environmental conditions. Therefore, it is of great interest to test if FB is also regulated by a sugar signal when plants are exposed to other abiotic stress conditions (e.g., low temperatures). In this context, comparative gene expression analysis (Fig. 4 and 5) also unveiled that other cellular processes and pathways, such as the biogenesis of ribosomes, which function as hubs for plant acclimation (Molina Garcia-Molina et al., 2020), were affected when TP export is perturbed. Hence, the importance of TP serving as metabolic retrograde signal (Pfannschmidt, 2010) and the connected cytosolic sugar biosynthesis and signalling pathways for HL acclimation should be analyzed in more detail in the future.

## Methods

### Genotypes and growth conditions

If not otherwise stated, *Arabidopsis thaliana* wild-type and mutant plants were grown on soil in short-day (SD, 10h light) at 100 μmol photons m^−2^ s^−1^ at 22°C. Genotypes used in this study are listed in Table S8. Homozygous *mpk6-2 and mpk6-3* were obtained from K.-J. Dietz (University Bielefeld, Ger), *gpt2-1, adg1-1 tpt2-1, xpt-1, xpt-2* and *gin2-1* from R. Häusler (University of Cologne, Ger) and *jassy* from S. Schwenkert (LMU Munich, Ger). Mutant lines overexpressing TREHALOSE 6-PHOSPHATASE form E. coli (otsB) were provided by A. Wiese-Klinkenberg (FZ Juelich) and *snrk1α1-3* amiRNAi *KIN11* by W. Dröge-Laser (University of Würzburg). T-DNA insertion mutants *pglct-2* and *tpt-2* and the EMS mutant *adg1-1* were obtained from the NASC seed stock centre. Gene knockout was confirmed by qPCR analysis or lack of starch in *adg1-1*. To induce the knockdown of *KIN11, snrk1α1-3* amiRNAi *KIN11* lines were grown for three weeks in SD and were sprayed without or with 20 μM ß-estradiol in water supplemented with 0.05% Tween-20 for another seven days.

### Treatment of plants

Standard high light shift experiments were carried out with 18-21d old plants exposed to 500 μmol photons m^−2^ s^−1^ for the indicated time-periods under constant temperature (22°C) in a Conviron Adaptis A/GEN1000 growth chamber equipped with a white LED light source. Plants were either shifted at the end of the night or 2 h after the onset of light. For young plants, samples contained leaves of approximately ten seedlings. For HL shift experiments with *snrk1α1-3* amiRNAi *KIN11*, two rosette leaves of four-week-old plants were harvested per sample, and each sample contained leaves from individual plants.

For high light shift experiments in normal (400ppm) or low (50ppm) CO_2_ conditions (21% oxygen), rosette leaves were mounted on a LICOR LI-6400XT device (LiCor, Lincoln, Nebraska USA) and incubated at 500 μmol photons m^−2^ s^−1^ (10% blue and 90% red LED light) for 4 h. The flow rate was set to 300μmol s^−1^ and block temperature to 22°C and relative humidity of 60%. After four hours, one leaf disc (2 cm^−2^) was punched out from the treated area of the leaf and directly frozen in liquid nitrogen. Six leaves from individual rosette plants were analyzed.

Sugar feeding experiments were performed by incubating plants in 20 mM TRIS buffer (pH 7.4) supplemented with 5% (w/v) sucrose.

For AZD treatment, seeds were surface sterilized by incubation in 70 % ethanol (plus 0.05 % Triton-X100) for 10 min. Subsequently, seeds were washed with 70 % ethanol, followed by a washing step in 100 % ethanol. After decanting the ethanol, seeds were dried and plated on 0.5x MS (4.4 g/l (w/v) MS including vitamins, 0.5 g/l (w/v) MES, 0.8 % agar, pH 5.7). MS plates were supplemented with 1μM AZD8055 (MedChemExpress, HY-10422). After stratification at 4°C for 2 d, plants were grown in SD conditions for 10d before the HL shift.

### Detection of reactive oxygen species

Superoxide radical accumulation was analyzed using Nitro blue tetrazolium chloride (NBT, Sigma-Aldrich, 93862). Leaves of a half rosette of around four weeks old plants were incubated in NBT staining solution (25 mM HEPES/KOH, 1 mg/ml NBT, pH 7.5). After 15-30 min vacuum infiltration, leaves were further incubated for 2h at RT in the dark. Hydrogen peroxide content was analysed using 3,3’-Diaminobenzidine (DAB, Merck-Millipore, D8001). The DAB staining solution (20 mM TRIS/Acetate, 1 mg/ml DAB (Sigma-Aldrich), pH 5) was prepared 1 h before use. After vacuum infiltration, leaves were kept in darkness 24h at RT. Chl was destained with 80 % (v/v) ethanol for 20 min at 80 °C in a water bath. Hydrogen peroxide contents were also analyzed using 2′,7′-dichlorofluorescein diacetate (H_2_DCF-DA, Sigma-Aldrich, 35845). To this end, leaves were incubated in 20 mM TRIS/HCl (pH 7.4) with or without 10 μM H_2_DCF-DA. After 15-30 min vacuum infiltration at RT, leaves were left in the solution for another 0.5-2h at RT in the dark. For the analysis, leaf discs were cut, and the fluorescence signal was detected using a ZEISS LSM 800 confocal laser-scanning microscope (63x magnification) at an excitation wavelength of 488 nm. The DCF signal was detected with an emission wavelength of 500-575 nm. Chlorophyll fluorescence emission was recorded between 650-680nm. The settings for analysis were the same for all analyzed samples.

### Quantification of starch

Starch was extracted from dried or frozen leaf material using 80 % (v/v) ethanol and incubation at 80 °C for 30 min. After centrifugation (10 min, 13000 rpm, RT), the soluble carbohydrate-containing supernatants were transferred to new tubes and prepared for sugar analysis (see below). The starch-containing pellet was resuspended in 750 μl of 0.5 M NaOH and incubated for 30 min at 95 °C before adding 750 μl of 1 M CH3COOH. Starch was digested by mixing 100 μl of the starch suspension with 100 μl of amyloglucosidase solution (1 mg/ml in 200 mM CH_3_COOH and 100 mM NaOH) for 2h shaking at 1050 rpm at 55 °C. Then, 100 μl of a 1:5, 1:10 or 1:20 dilution of the starch digestion diluted in H_2_O_dd_ were mixed with 200 μl of glucose oxidase (GlcOx) reagent (1 mg glucose oxidase, 1.5 mg horseradish peroxidase, 5 mg dianisidine/HCl per 50 ml 0.5 M TRIS/HCl (pH 7.0), 40 % (v/v) glycerol). For the glucose standard curve, 0-1 mM glucose was mixed with 200 μl of the GlcOx. Standard curve and samples were incubated at 30°C for 30 min before the reaction was stopped by the addition of 400 μl of 5 M HCl. After short centrifugation (30 sec, RT), the absorbance of the samples at 540 nm was determined using a 96-well plate reader (SpectraMax® M2 microplate reader (Molecular Devices LLC., USA)). The starch amount was determined as glucose equivalents, and glucose content was calculated using the standard curve.

### Quantification of soluble sugars

Concentrations of free sugars sucrose, glucose and fructose were quantified photometrically as previously described Atanasov et al., 2020. Supernatants from the first step of the starch extraction (see above) were collected and dried in a speed vacuum concentrator. Sugars were extracted from dried pellets in H_2_O_dd_ by constantly shaking with 500 rpm at 22°C. For sucrose quantification, sample extracts were incubated for 10 min with 30% KOH at 95°C, followed by incubation for 30 min at 40°C with an anthrone reagent (0.14% w/v anthrone in 14.6 M H_2_SO_4_). Together with a standard calibration curve, sample absorbance was determined at 620 nm. Glucose concentrations were determined from extracts in a coupled hexokinase/glucose-6-phosphate dehydrogenase assay resulting in NADPH + H^+^ formation, detectable at 340 nm. Following glucose quantification, phosphoglucose isomerase was added to the mixture to determine fructose concentrations. The absolute amount was calculated using standard calibration curves.

### Anthocyanin extraction and quantification

Anthocyanins were extracted from the ground and frozen leaf material using 1ml of anthocyanin extraction buffer (18% 1-propanol, 1% HCl in water). Homogenates were incubated for 2 h at RT in darkness. After centrifugation for 10 minutes at RT (13000 rpm), supernatants were transferred to cuvettes, and the absorption at 547nm, 650nm and 720nm was recorded. The absorption of anthocyanins was calculated using the following formula: (A_537_-A_720_)-0.25x(A_650_-A_720_). Absorption values were normalized to either g fresh weight (fw) or dry weight (dw).

### Protein extraction and western blot analysis

Total leaf proteins were extracted from ground and frozen leaf material using protein extraction buffer (56 mM Na_2_CO_3_, 56 mM DTT, 2% (w/v) SDS, 12% (w/v) sucrose, 2 mM EDTA). After resuspending the powder, samples were incubated at 90°C for 10 min, and protein extracts were obtained by subsequent centrifugation (10 min, 13.000 rpm, RT). Protein extracts were separated on 12% Polyacrylamide SDS-gels and blotted onto nitrocellulose membrane. After blocking for 1 h in 4 % milk solution in TBS-T (50 mM TRIS/HCl, 150 mM NaCl, pH 7.5; 0.1 % (v/v) Tween-20), membranes were incubated overnight at 4°C with the primary antibody in 1 % milk solution in TBS (phospho-AMPK, 1:2000, Cell Signaling Technology, #2535). The next day, membranes were washed and incubated with the secondary antibody for 2 h at RT (goat anti-rabbit IgG coupled with HRP, 1:10.000, 1 % milk solution, TBS). Signals were detected using Clarity Western ECL substrate (Bio-Rad, Ger) and an ECL Chemostar CCD-camera (Intas, Ger).

### RNA extraction and qPCR analysis

Whole leaf RNA was extracted using previously published protocol (Oñate-Sánchez and Vicente-Carbajosa, 2008). In brief, frozen and ground leaf material was resuspended in 300μl cell lysis buffer (2 % (w/v) SDS, 68 mM sodium citrate, 132 mM citric acid, 1 mM EDTA). After the addition of 100μl DNA/protein precipitation solution (4 M NaCl, 16 mM sodium citrate, 32 mM citric acid), samples were vortexed and incubated on ice for 10 min. Subsequently, samples were centrifuged at 13000rpm (4°C) for 10 min, and 300μl of the supernatant were used to precipitate the RNA with 300μl Isopropanol. After precipitation (5 min, 13.000 rpm, RT), pellets were washed with 800μl 75% ethanol, centrifuged again and dried. The RNA was resuspended in 20-25μl of H_2_O_dd_ (RNAse free) and stored at −80°C until further use.

For qPCR analysis, 1-2μg of DNaseI (Thermofisher) treated RNA was transcribed into cDNA using RevertAid reverse transcriptase (Thermofisher) according to manufacturer protocol. qPCR analysis was carried out in a CFX96-C1000 96-well plate thermocycler (Bio-Rad) and ChamQ Universal SYBR® qPCR Master Mix (Absource) in 6μl reactions containing 1μl of diluted (1:5) cDNA. Relative gene expression was calculated using the 2^−ΔΔ(C(t))^ method and *SAND* (AT2G28390) as the reference gene. Primers used for qPCR analysis are listed in Table S9.

For RNA-sequencing, RNA was treated with DNase I and purified using phenol/chloroform/isoamyl alcohol precipitation protocol. After washing with 75% ethanol (v/v), RNA was dried and resuspended in RNase-free water.

### RNA-sequencing and data analysis

RNA-seq was performed by Novogene (UK/China). Three biological replicates for Col-0 and *tpt-2* after 9 h and 18 h of HL treatment were analyzed. Messenger RNA was purified from total RNA using poly-T oligo-attached magnetic beads. After fragmentation, the first-strand cDNA was synthesized using random hexamer primers, followed by the second strand cDNA synthesis. The library was tested with Qubit and real-time PCR for quantification and bioanalyzer for size distribution detection. Quantified libraries were sequenced on Illumina platforms (HiSeq4000), according to effective library concentration and data amount (4GB raw reads). Raw data of fastq format were processed through in-house perl scripts. This step obtained clean data (clean reads) by removing reads containing adapter, poly-N, and low quality. At the same time, Q20, Q30 and GC content of the clean data were calculated.

Paired-end clean reads were aligned to the reference genome using Hisat2 v2.0.5. Reference genome and gene model annotation files of the TAIR10 release were used. featureCounts v1.5.0-p3 was used to count the reads numbers mapped to each gene. FPKM (Fragments Per Kilobase of transcript sequence per Millions of base pairs sequenced) of each gene was calculated based on the length of the gene and reads count mapped to this gene. Differential gene expression analysis (three biological replicates per condition) was performed using the DESeq2 R package (1.20.0). The resulting p-values were adjusted using Benjamini and Hochberg’s approach for controlling the false discovery rate. Post-sequencing analysis of data was performed using Galaxy webserver (Galaxy Europe, https://usegalaxy.eu/), a tool for the preparation of venn diagrams (https://bioinfogp.cnb.csic.es/tools/venny/), and R studio (heatmap package: ‘pheatmap’ and correlation analysis: ‘corrplot’). Gene ontology (GO) term analysis was performed using GO TERM FINDER with settings for Arabidopsis thaliana (https://go.princeton.edu/cgi-bin/GOTermFinder). Own transcriptome data was compared to expression data published in Huang et al., 2019, Baena-Gonzalez et al., 2007 and Garcia-Molina et al., 2020.

## Data availability

Transcriptome data was deposited at GEO (https://www.ncbi.nlm.nih.gov/geo/info/update.html) under the record GSE196053.

## Author Contributions

MEZ, GEA, AK, KJ, TR, CK, TN, ASR performed the experiments and analyzed the data. A.S.R. designed, conceived and supervised the study and wrote the article with the support of all authors. A.S.R. and T.N. acquired funding.

## Acknowledgement

We want to acknowledge the generous gifts of seeds obtained from the following colleagues: K.-J. Dietz (*mpk6*), W. Dröge-Laser (*snrk1α1-3* amiRNAi *KIN11*), R. Häusler (*gpt-2, adg tpt, xpt, gin2-1*, A. Wiese-Klinkenberg (*otsB*) and S. Schwenkert (*jassy*). We also thank R. Häusler for discussion on the project.

## Figures

**Figure S1:**
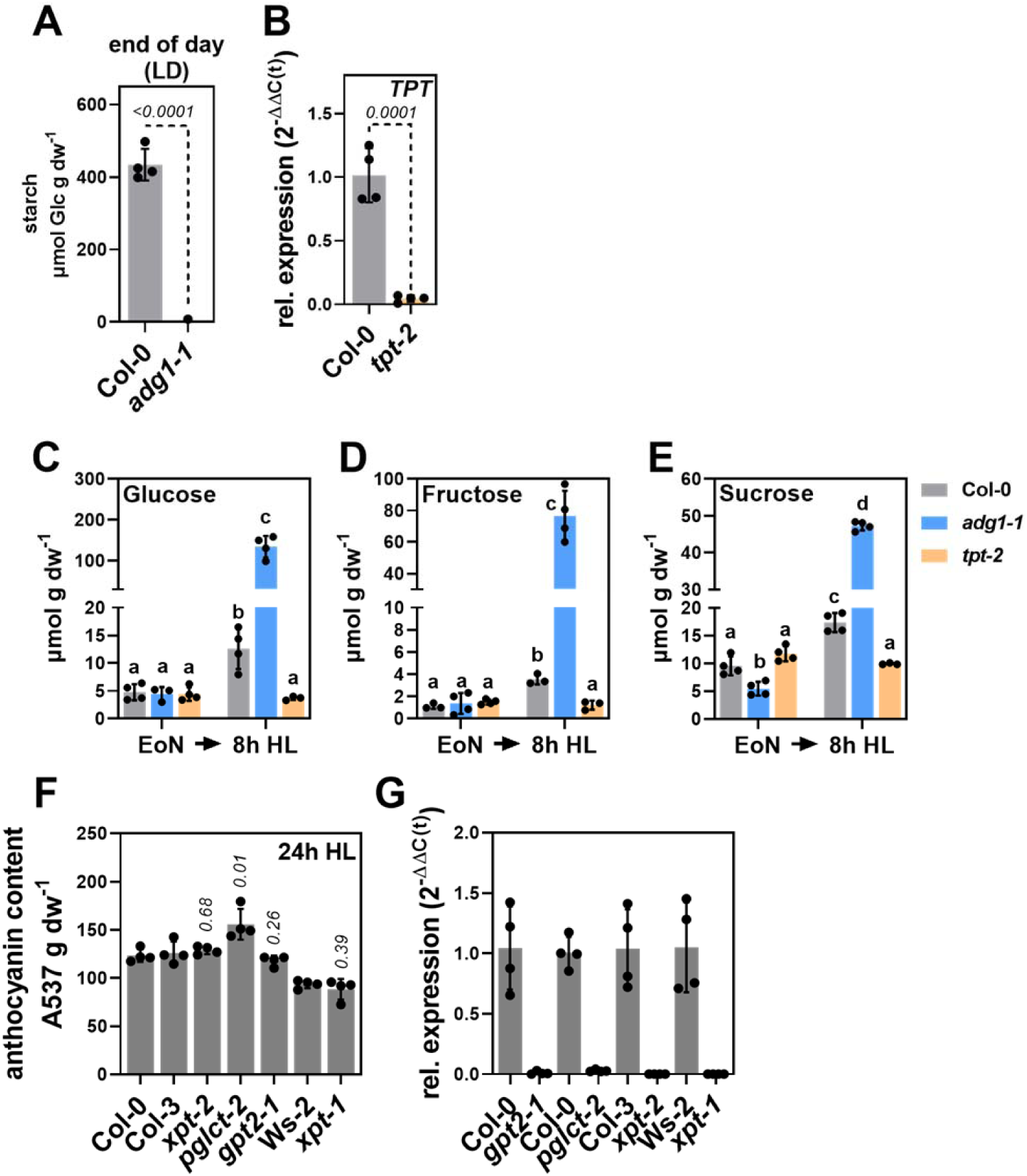
(**A**) Analysis of starch content in the *adg1-1* mutant at the end of day (long-day), (**B**) *TPT* expression in Col-0 and *tpt-2* mutant, (**C**) glucose, (**D**) fructose, (**E**) sucrose content in Col-0, *adg1-1*, and *tpt-2*. Plants grown in SD were analyzed at the end of the night (EoN) and after 8h HL shift. (**F**) Anthocyanin content in mutants for plastid-localized transporters after 24h HL treatment. The *xpt-1* mutant was in Wassilewskija (Ws-2) and *xpt-2* in Col-3 background. (**G**) Confirmation of gene knockout in mutants shown in (F). For (A) Relative gene expression was calculated using the 2^−ΔΔ(C(t))^ method and *SAND* as reference gene relative to Col-0. For (G) the control was the respective WT background. For (A), (B) and (F), statistical significance between wild-type and mutant(s) was analyzed using student’s t-test. Values are mean ±SD (n=4) and the *p*-values are shown. For (C-E) statistical significance between genotypes was analyzed by two-way ANOVA (Tukey’s multiple comparisons test) analysis (n≥3) and significance groups are indicated by letters (*p*<0.01).

**Figure S2:**
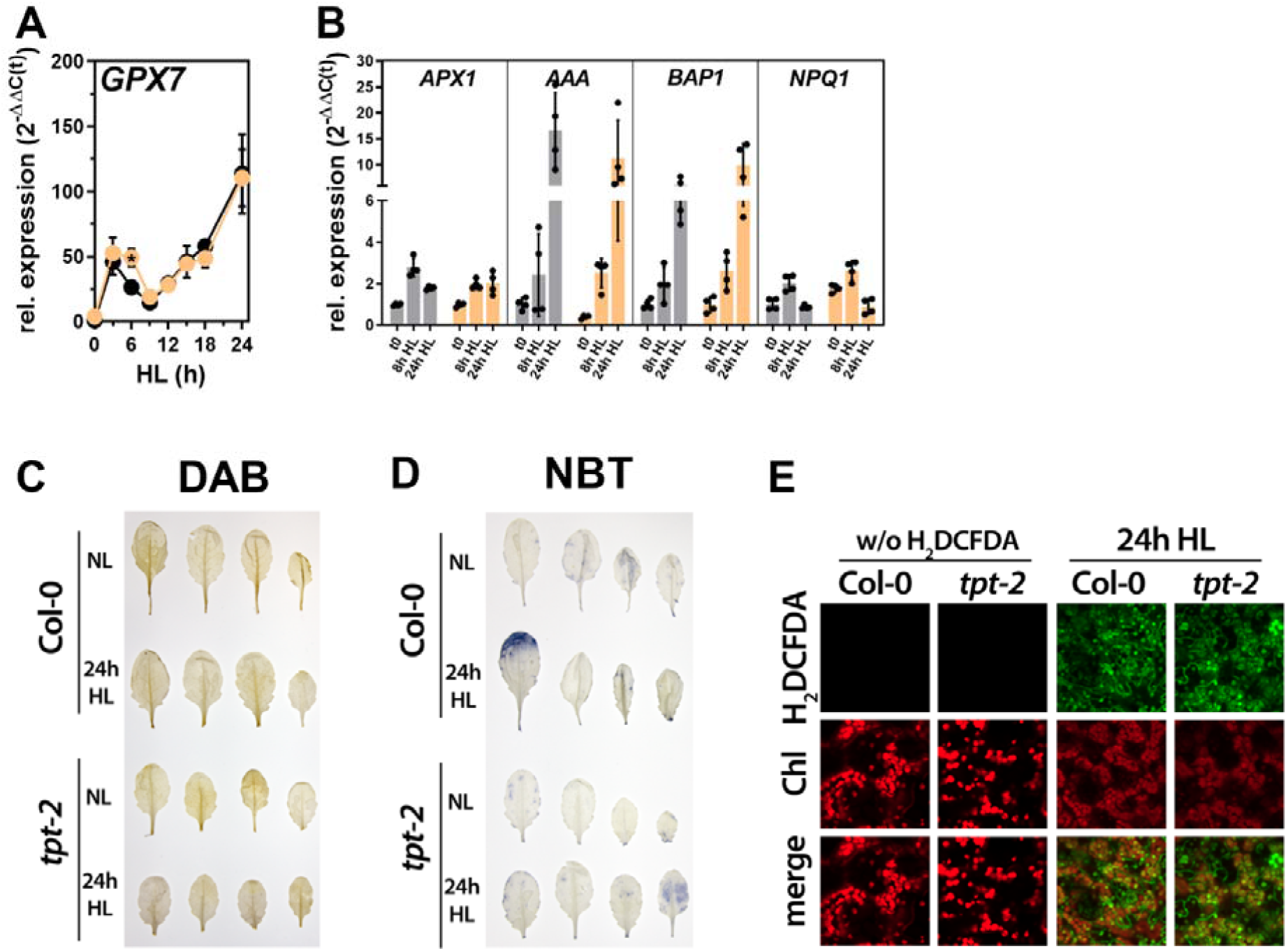
The *tpt-2* mutant showed a WT-like ROS response during HL shifts. (**A**) Expression of *GLUTATHIONE PEROXIDASE 7 (GPX7*, AT4G31870) during a 24h HL shift experiment (compare Fig. 2), (**B**) *ASCORBATE PEROXIDASE1 (APX1*, AT1G07890*), AAA-ATPase (AAA*, AT3G28580*), BON ASSOCIATION PROTEIN 1 (BAP1*, AT3G61190*) and VIOLAXANTHIN DEEPOXIDASE (NPQ1*, AT1G08550) in Col-0 and *tpt-2* at t0 and the indicated time-points after the HL shift. Changes in gene expression were calculated using the 2^−ΔΔ(C(t))^ method relative to Col-0 before the HL shift (t0) and *SAND* as reference gene. Values are mean ±SD (n=4). (**C**) 3,3’-Diaminobenzidine (DAB), (**D**) Nitro blue tetrazolium chloride (NBT) and (**E**) 2′,7′-dichlorofluorescein diacetate (H_2_DCF-DA) staining of leaves from normal light (NL) conditions and after 24h of continuous HL. Signals for chlorophyll autofluorescence (Chl, red, emission 650-680 nm) and DCF (green, emission 500-575 nm) were recorded using a confocal laser scanning microscope and 488 nm excitation wavelength. Further details are given in the materials and methods section. Leaf samples were analyzed with the same settings.

**Figure S3:**
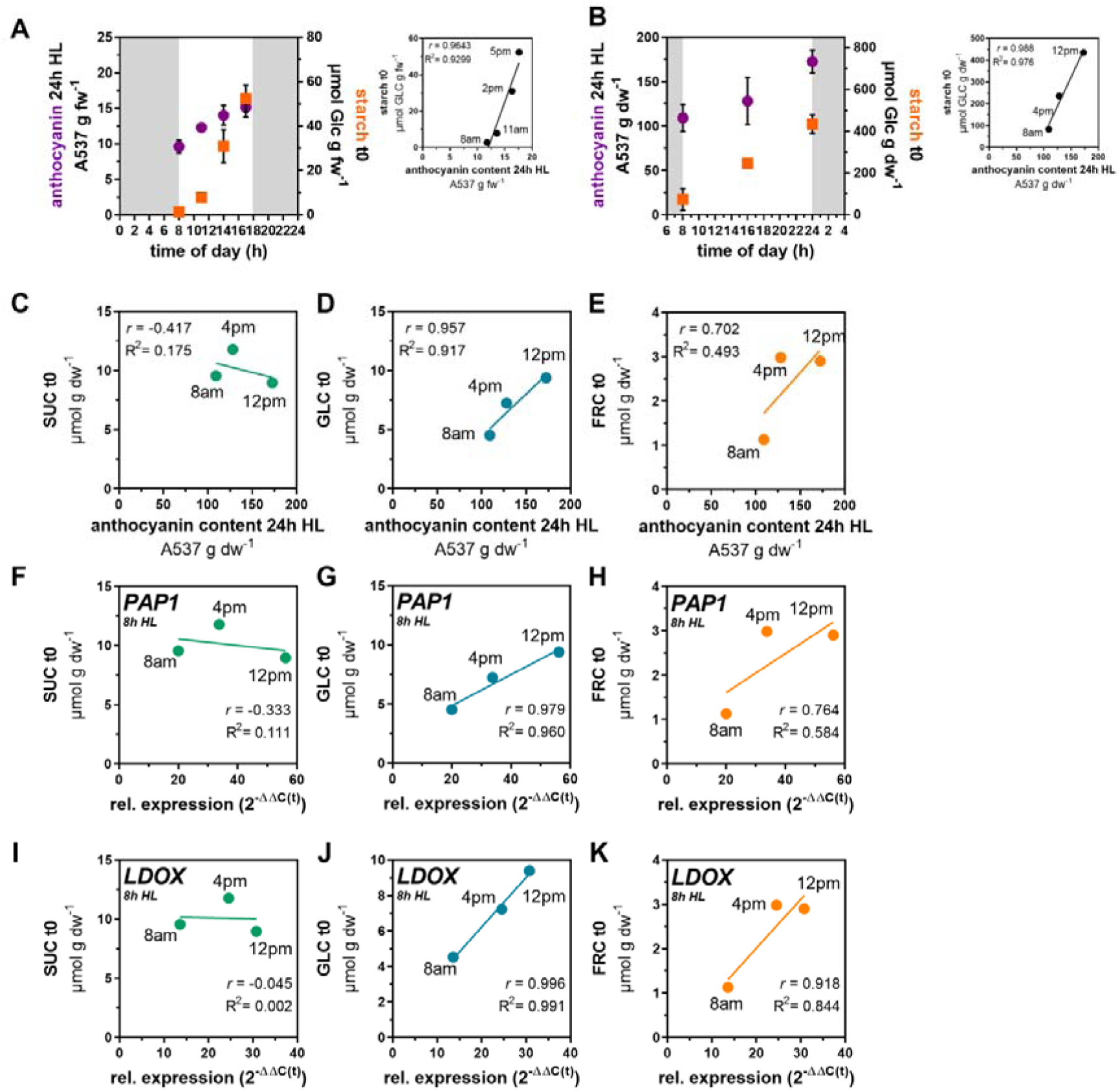
Induction of flavonoid biosynthesis in HL correlated with starch and sugar contents before the HL shift. (**A**) Short-day and (**B**) long-day grown *Arabidopsis* wild-type were subjected to 24h HL treatment at different time points during the day. Samples for starch analysis (orange) were harvested before the HL shift. After 24 h of continuous HL treatment, anthocyanin contents (purple) were quantified. Correlation analysis of starch contents before (t0) and anthocyanin contents after 24 h HL treatment are shown to the right of each graph. Values in (A) represent the mean ±SEM for n=8 samples from three independent experiments. In (B), the mean ±SD for n≥3 are shown. (**C-E**) Correlation analysis of anthocyanin content after 24 h HL treatment and (C) sucrose (SUC), (D) glucose (GLC) and (E) fructose (FRC) contents before the HL shift. To reduce the complexity of the graphs, only the mean values of n≥3 samples are shown. (**F-K**) Relative expression of *PAP1* (**F-H**) and *LDOX* (**I-K**) in Col-0 after 8h of HL treatment was correlated with SUC (F and I), GLC (G and J) and FRC (H and K) contents before the HL shift (t0). Changes in gene expression were calculated relative to the expression values at 8 am prior to the HL shift. Gene expression was calculated using the 2^−ΔΔ(C(t))^ method and *SAND* as reference gene. To reduce the complexity of the graphs, only the mean values of n≥3 samples are shown. Pearson correlation coefficient (*r*) and the linear regression correlation coefficient (R^2^) are shown.

**Figure S4:**
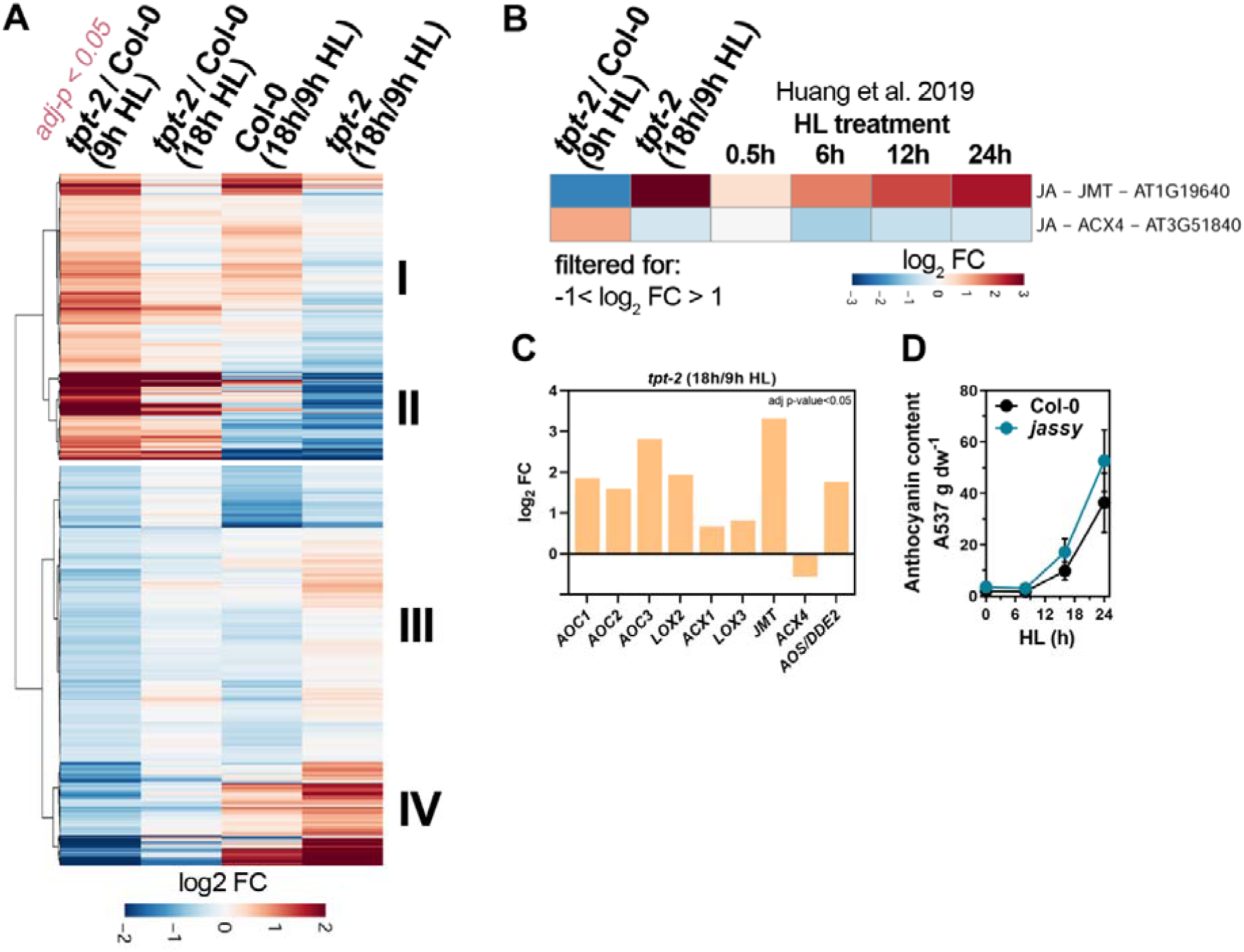
**(A)** RNA-seq analysis of significantly differentially expressed genes (DEGs) in *tpt-2* and Col-0 at different time-points of the HL kinetic. Transcripts were filtered for DEGs (adjusted p-value<0.05) in *tpt-2* relative to Col-0 at 9 h of the HL treatment, and changes of expression in the indicated comparisons are shown. Cluster I and II contain moderately to highly induced transcripts in *tpt-2*/Col-0 at 9 h HL whose expression is largely repressed after 18h of HL treatment in *tpt-2* relative to 9 h HL treatment (last column). Cluster III and IV encompass transcripts that were moderate to strongly downregulated in *tpt-2*/Col-0 at 9 h HL but were induced in *tpt-2* after 18h of HL treatment relative to 9 h HL. (**B**) Relative expression changes for *JASMONIC ACID CARBOXYL METHYLTRANSFERASE (JMT)* and *ACYL-COA OXIDASE 4 (ACX4)*. The list of DEGs in *tpt-2*/Col-0 (9 h HL) was filtered (−1<log_2_ FC>1, adj-p<0.05) for hormone biosynthesis genes and downstream targets of hormone signalling and only *JMT* and *ACX4* were found to be deregulated in *tpt-2* at 9h HL (Huang et al. 2019, see also Table S3). Changes in gene expression are given as log_2_ fold change (FC) relative to the control and hierarchical row clustering (Euclidean distance) using ward.D method was applied. (**C**) Expression of DEG involved in jasmonate biosynthesis in *tpt-2* 18 h/9 h HL (adjusted p-value<0.05). *JASMONIC ACID CARBOXYL METHYLTRANSFERASE (JMT)* and *ACYL-COA OXIDASE (ACX1* and *4), LIPOXYGENASE* (*LOX2* and *3*), *ALLENE OXIDE SYNTHASE* (*AOS*) and *ALLENE OXIDE CYCLASE* (*AOC1-3*). (**D**) Accumulation of anthocyanins in Col-0 (black) and jasmonate-deficient *jassy* mutant (blue) during a 24h HL shift. Values are given as mean ±SD for n≥3 samples.

**Figure S5:**
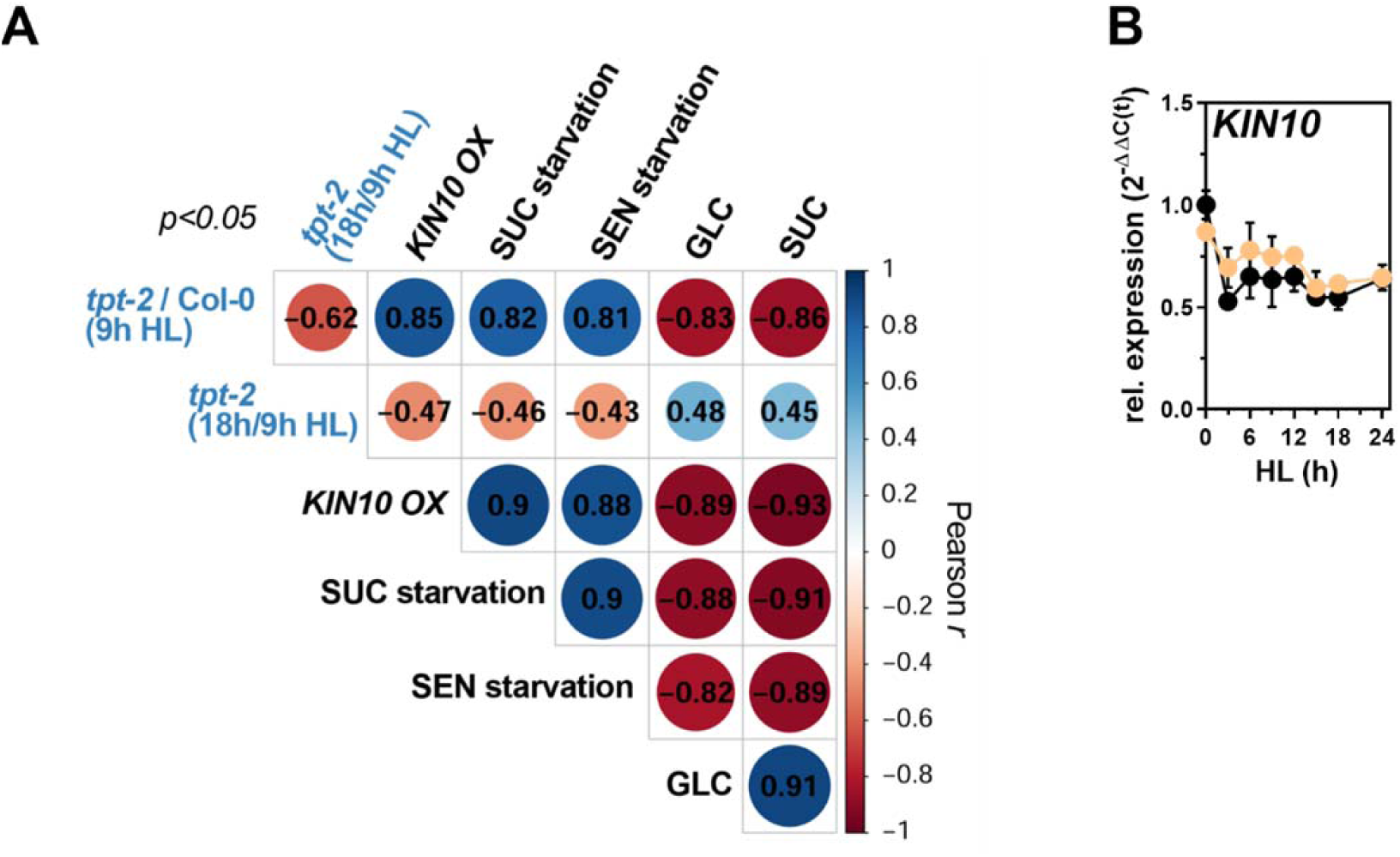
(**A**) Correlation plot of transcriptomes shown in Figure 5B. The size and colors represent the Pearson correlation coefficient (*r*, depicted inside the circles). Note that the changes in the *tpt-2* relative to Col-0 at 9 h HL were positively correlated with transcriptome changes induced by *KIN10* overexpression in protoplasts and starvation conditions but negatively correlated with gene expression changes stimulated by glucose (GLC) and sucrose (SUC) feeding. In contrast, compared to 9 h HL prolonged HL treatment induced changes in gene expression in *tpt-2*, leading to a negative correlation with *KIN10 OX* and starvation but a positive correlation with sugar feeding. (**B**) Expression of the catalytic SnRK1 subunit *KIN10* in Col-0 (black) and *tpt-2* (orange) through a HL shift kinetic (compare Figure 2).

**Figure S6:**
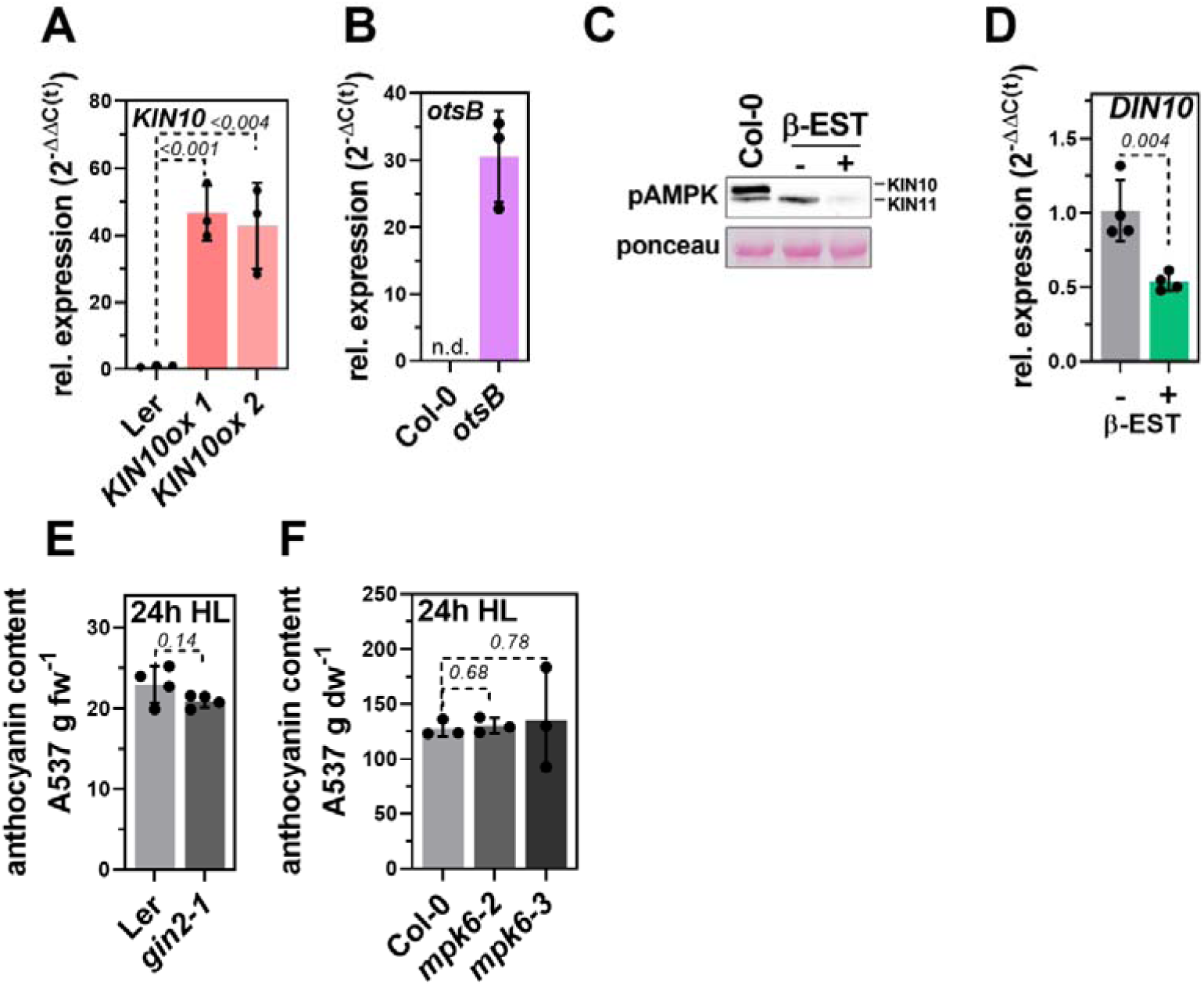
Confirmation of (**A**) *KIN10* overexpression in two independent *KIN10ox* lines, and **(B)** *otsB* expression in *otsB*. (**C**) Western-blot confirming the knockout of KIN10/SnRK*α*1 and knockdown of KIN11/SnRK*α*2 in *snrk1α1-3* amiRNAi *KIN11* induced by ß-EST application. The content of both catalytic SnRK1 subunits was analzed using an antibody raised against the phosphorylated T-loop (T172) of human AMP-activated protein kinase (pAMPK) recognizing also phosphorylated SnRK1α1^T175^/SnRK1α2^T176^. (**D**) *DIN10* expression in *snrk1a1-3* amiRNAi *KIN11* at the end of a night (14h dark). Gene expression was calculated using the 2^−ΔΔ(C(t))^ (for (A and C)) and 2^−ΔΔ(C(t))^ (for B) method relative to the expression in the WT/untreated control (for A and B), and *SAND* as reference gene. Statistical significance between Ler and *KIN10ox* was analyzed using student’s t-test. Values are mean ±SD (n≥3) and the *p*-values are shown. n.d., not detectable. (**E**) Anthocyanin content in Ler and *gin2-1* after 24h HL (n=4). (**F**) Anthocyanin content in Col-0, *mpk6-2* and *mpk6-3* mutants after 24 h HL (n=3). For (D) and (E) Statistical significance between WT and mutant(s) was analyzed using student’s t-test and the p-values are shown.

